# Colitis-associated intestinal microbiota regulates brain glycine and host behavior in mice

**DOI:** 10.1101/2022.03.07.483210

**Authors:** Maryana V. Morozova, Mariya A. Borisova, Olga A. Snytnikova, Ksneniya M. Achasova, Ekaterina A. Litvinova, Yuri P. Tsentalovich, Elena N. Kozhevnikova

## Abstract

Inflammatory bowel diseases (IBD) are chronic and relapsing inflammatory disorders of the gastrointestinal tract with complex etiology and no strategies for complete cure. IBD are often complicated by mental disorders like anxiety and depression, indicating substantial shifts in the gut-brain axis. However, the mechanisms connecting IBD to mental diseases are still under debate. Here we use *Muc2* mutant mouse model of chronic colitis to uncouple the effects of the intestinal microbiota on host behavior from chronic inflammation in the gut. *Muc2* mutant male mice exhibit high exploratory activity, reduced anxiety-related behaviors, impaired sensorimotor gating, and altered social preference towards males and females. Microbial transfer to wild-type mice via littermate co-housing shows that colitis-associated microbiota rather than inflammation *per se* defines behavioral features in *Muc2* colitis model. Metagenomic profiling and combination of antibiotic treatments revealed that bacterial species *Akkermansia muciniphila* is associated with the behavioral phenotype in mutants, and that its intestinal abundance correlates with social preference towards males. Metabolomic analysis together with pharmacological inhibition of Gly and NMDA receptors helped us to determine that brain glycine is responsible for the behavioral phenotype in *Muc2* mice. Blood and brain metabolic profiles suggest that microbiota-dependent changes in choline metabolism might be involved in regulation of central glycine neurotransmission. Taken together, our data demonstrates that colitis-associated microbiota controls anxiety, sensorimotor gating and social behavior via metabolic regulation of the brain glycinergic system, providing new venues to combat neurological complications of IBD.

## Introduction

IBD are gastrointestinal (GI) disorders involving inflammation, ulcerations, epithelial barrier dysfunction, diarrhea, weight loss, anemia and predisposition to colorectal cancer (Baumgart and Sandborn, 2007; Feagins et al., 2009; Gasche, 2000; Romano et al., 2017). IBD onset is on average at 15-35 years of age, and it remains a chronic and relapsing disorder throughout patient’s life. The etiology of IBD is under debate, with genetic predisposition, environmental exposures and diet emerging as the main risk factors (Korzenik, 2005; Neuman and Nanau, 2012; Williams et al., 2002). Human studies support the role of heritable aspects in IBD, since first-degree relatives of IBD patients have significantly higher risk to develop this illness than the background population (Kevans et al., 2016; Park et al., 2006; Santos et al., 2018). Genome-wide screens revealed about 200 genetic loci linked to IBD, many of which are immune system components (Jostins et al., 2012; De Lange et al., 2017; Liu et al., 2015). Animal models demonstrate that mutations in some of these genes predispose to intestinal inflammation upon environmental triggers, but may not be sufficient to induce colitis (Kullberg et al., 1998). These data suggest that IBD arise on the interface of mucosal and intestinal environment, both being indispensable for the disease progression (MacDonald et al., 2011). The main component of the heritable intestinal environment is microbiota, which is in a mutual regulatory relationship with mucosal immune system (Cerf-Bensussan and Gaboriau-Routhiau, 2010; Thaiss et al., 2016; Zheng et al., 2020). Indeed, multiple reports describe a significant association of intestinal inflammation with distinct microbiome signatures that might induce IBD as well as result from it (Chassaing and Darfeuillemichaud, 2011; Iljazovic et al., 2021; Kostic et al., 2014). In the experimental models of colitis, transplants of fecal microbiota or inoculation with probiotic bacterial species alleviated the severity of inflammation (Ding et al., 2019; Guslandi et al., 2000; Thomas, 2016). Therefore, intestinal microbes can beneficially regulate pathological aspects of IBD.

It has been noticed that IBD are often accompanied by mental disorders including anxiety, depression, bipolar disorder, insomnia, lack of motivation, and others (Conley et al., 2021; Neuendorf et al., 2016). About 35-40% of IBD patients suffer from anxiety and about 22-25% - from depression, indicating a strong interaction between intestinal inflammation and the central nervous system (CNS) (Moulton et al., 2019; Neuendorf et al., 2016; Selinger and Bannaga, 2015). Particularly, microbial metabolism substantially impacts the overall metabolomic profile of the host, including amino acids, short chain fatty acids, lipids and intermediary metabolites (Abdul Rahim et al., 2019; Fan and Pedersen, 2021; Xiao et al., 2020). These microbiota-associated molecules can potentially affect host neurophysiology and behavior serving as precursors for neurotransmitter biosynthesis and by regulating neuro- and axonogenesis (Huang and Wu, 2021; Sampson and Mazmanian, 2015; Strandwitz, 2018; Vuong et al., 2017). However, the mechanisms underlying mental disorders during IBD are still unclear due to the complexity of this disease. In order to understand the interrelation between chronic colitis, microbiota and CNS function, we utilized *Mucin 2* (*Muc2*) knockout mice as a well-described model of intestinal inflammation (Lu et al., 2011; Velcich et al., 2002a). This mutant animals lack the major secreted mucin in the intestine and reconstitute key features of chronic colitis (Bergstrom et al., 2010; Johansson et al., 2008; Van der Sluis et al., 2006). Here we show that intestinal microbiota regulates brain glycine and controls anxiety-related and social behaviors in *Muc2* mutant mice. These behavioral traits were associated with elevated intestinal *Akkermansia muciniphila* and were transferrable along with the intestinal microbiota to wild-type animals. We propose a mechanism, by which *Akkermansia muciniphila* affects host betaine-glycine metabolism leading to activation of glycine-dependent neurotransmission. Therefore, glycine emerges as a potential mediator of the gut-brain axis regulated by intestinal microbiota.

## Results

### Behavioral phenotyping of Muc2^−/−^ male mice reveals high novelty-induced activity, reduced anxiety-related behavior, impaired startle reflex and abnormal social behavior

First, we used behavioral phenotyping to investigate whether chronic inflammation in *Muc2* knockout model has a physiologically relevant effect on CNS. We evaluated behavioral pattern in the home cage environment using Phenomaster technology (Figure 1A). Among parameters tested, *Muc2^−/−^* mice showed a reduced motor activity and an increased water consumption rate as compared to C57BL/6. Most probably, increased drinking is a result of diarrhea in mutants. At the same time, locomotion was increased in a novel environment in *Muc2^−/−^* males as revealed by the open field test (Figure 1B). *Muc2*^−/−^ animals had increased horizontal activity at the level of a trend and significantly increased vertical activity (rearings). Moreover, *Muc2*^−/−^ mice spent significantly more time in the center of the arena indicating reduced anxiety. These results were supported by the light-dark test, where mutant animals showed an increased motor activity, more time and number of entries in the light compartment (Figure 1C). Some reduction in anxiety was also noted in the elevated plus-maze where *Muc2^−/−^* mice had an increased number of peeks down from the open arms (Figure S1). However, there were no changes in the distance and time in open arms. In the marble burying test, *Muc2*^−/−^ animals buried significantly fewer marbles as compared to C57BL/6 indicative of the reduced repetitive behavior (Figure 1D). Behavioral phenotyping revealed no depressive-like traits and a superior motor performance in *Muc2^−/−^* mice (Figure S1). We also found no impairment of cognitive functions, since there were no substantial differences in spontaneous alternation, long-term memory retention and active avoidance in mutants as compared to the control (Figure S1). Next, we studied sensorimotor gating in *Muc2* model by measuring prepulse inhibition (PPI), startle reflex and habituation to the acoustic signal. We found no difference in PPI between C57BL/6 and *Muc2*^−/−^ animals, however, mutant mice had significantly weaker startle reflex and diminished habituation to the sound (Figure 1E). Thus, colitis affected neuronal circuits responsible for sensorimotor gating in *Muc2^−/−^* mice.

**Figure 1.**
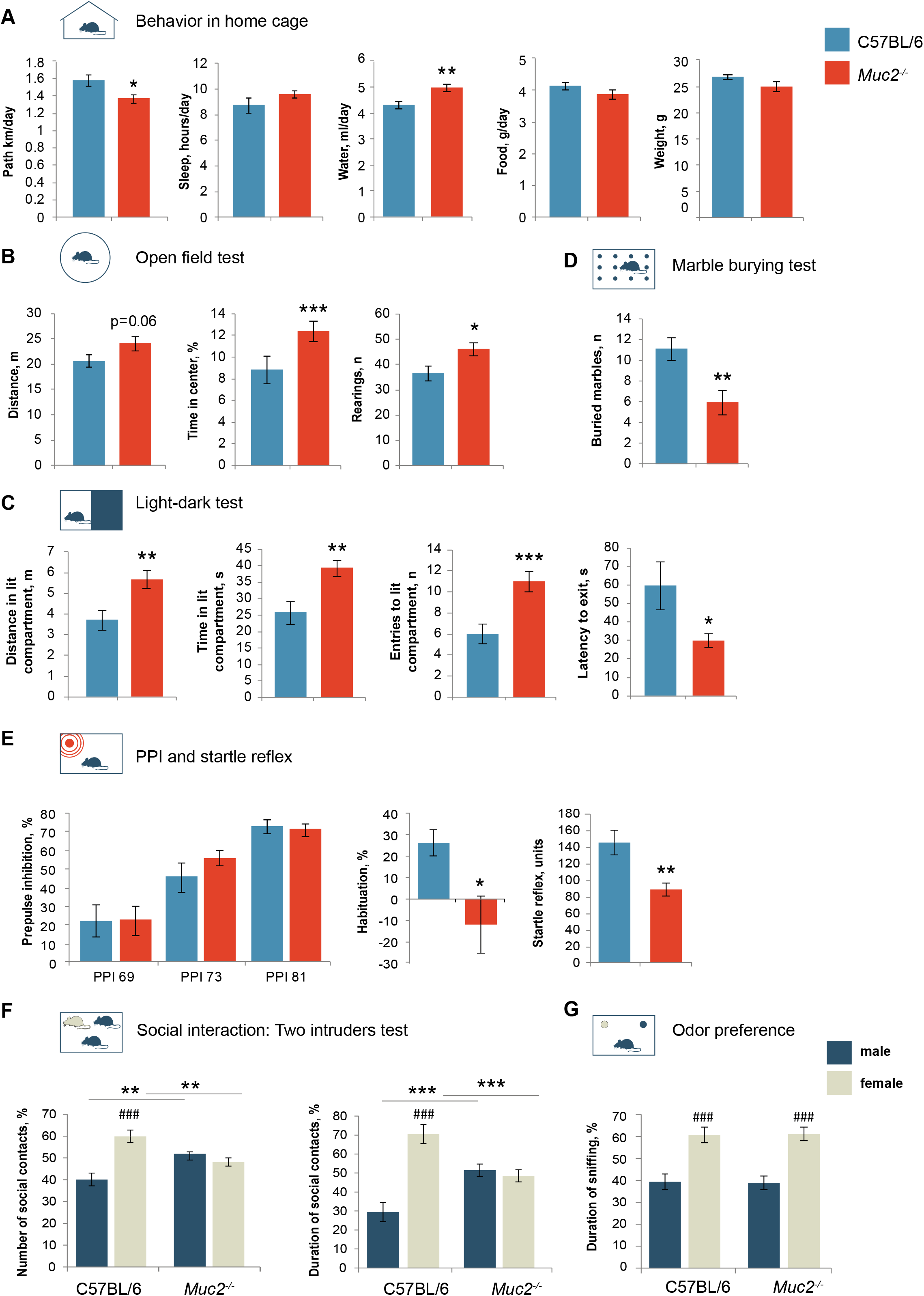
Behavioral traits of *Muc2^−/−^* animals. **(A)** Behavior in home cage (*n* = 9-12). Motor activity: *t* = 2.46, *p* = 0.024; water consumption rate: *t* = −3.59, *p* = 0.002, Student’s *t*-test. **(B)** Open field test (*n* = 18-20). Distance: *t* = −1.91, *p* = 0.06, Student’s *t*-test; rearings: *t* = −2.326, *p* = 0.026, Student’s *t*-test; time in the center: Z = −3.333, *p* < 0.001, Mann-Whitney *u* test *vs*. C57BL/6. (**C)** Light-dark test (*n* = 17-20). Distance: *t* = −3.067, *p* = 0.004; time: *t* = −3.08, *p* = 0.004; entries: *t* = −3.66, *p* < 0.001, Student’s *t*-test. **(D)** Marble burying test (*n* = 17-20). Number of buried marbles: Z = 2.93, *p* = 0.003; Mann-Whitney *u* test. **(E)** Habituation and startle reflex (n = 16-18). Startle reflex: *t* = 3.246, *p* = 0.003, Student’s *t*-test; habituation: Z = 2.17, *p* = 0.03; Mann-Whitney *u* test. **(F)** Two intruders test (*n* = 10-11). There was a statistically significant interaction between the resident genotype and the intruder gender (number of contacts: F(1, 38) = 21.990, *p* < 0.001; duration: F(1, 38) = 26.045, *p* < 0.001, two-way ANOVA). Number of contacts (C57BL/6, male *vs*. female): *p* < 0.001, Fisher’s LSD test; duration (C57BL/6, male *vs*. female): *p* < 0.001, Fisher’s LSD test. Number of contacts with a male/female intruder *vs*. C57BL/6: *p* < 0.01, Fisher’s LSD test; duration of contact with a male/female intruder *vs*. C57BL/6: *p* < 0.001, Fisher’s LSD test. **(G**) Odor preference test (*n* = 12-21). There was a statistically significant influence of the gender of animal whose odor was presented to the tested male F(1, 62) = 34.17 *p* < 0.001 two-way ANOVA. Duration of sniffing (male *vs*. female): *p* < 0.001, Fisher’s LSD test. * = *p* < 0.05, ** = *p* < 0.01, *** = *p* < 0.001, *vs*. C57BL/6. ### = *p* < 0.001, male *vs*. female.

Social preference test revealed that *Muc2* mutation did not result in autistic phenotype, since mutant mice preferred an animal to an unanimated object in a social preference test (Figure S1). However, in contrast to C57BL/6 animals, mutant males lack preference towards female intruders and tend to interact equally with a male and a female intruder in a two intruders paradigm (Figure 1F). Control males strongly preferred to contact a female than a male, whereas mutant mice demonstrated no preferences towards either. Moreover, *Muc2^−/−^* males interacted with a male intruder significantly more than did the control animals, revealing a substantial shift in social preference. These results were supported by the resident-intruder test with a male mouse, where *Muc2^−/−^* males made more contacts to the intruder, with more animals exhibiting aggression or even mating to males (Figure S2). At the same time, when a female mouse was used as an intruder, mutant males mated females at a lower rate than did the wild type controls (Figure S2). Likewise, in the two intruders test, mutant males were more likely to mate male intruders and attack females, without any behavioral discrimination between male and female intruders (Figure S2). In order to test whether *Muc2* mutation affects perception of socially-relevant olfactory signals we performed an odor preference test. By providing test animals with bedding from female and male cages we found that mutant males discriminated male and female odors at the same rate as their control counterparts (Figure 1G).

Altogether, the main behavioral traits characteristic to *Muc2* colitis model are elevated novelty-induced locomotion and exploratory activity, reduced anxiety-related behaviors, weakened startle response and lack of behavioral discrimination between male and female animals.

### Behavioral traits of Muc2^−/−^ mice are transferrable to the wild-type animals upon littermate co-housing

Our *Muc2* model behavioral data unexpectedly demonstrate enhancement of exploratory activity and reduction in anxiety, which is generally not common in chronic intestinal inflammation. Thus, we suggested that the observed phenotype isattributed to the colitis-associated microbiota rather than inflammation *per se*. As mucosal environment strongly affects intestinal flora, we performed littermate co-housing of *Muc2*^−/−^ and C57BL/6 animals in order to facilitate microbiota exchange and test its involvement in the behavioral phenotype. C57BL/6 were crossed to *Muc2*^−/−^ animals to generate heterozygous mutants, which were mated to obtain *Muc2*^+/+^ (wild-type littermates) test group (Figure 2A). These animals had no features of intestinal inflammation or any clinical signs of colitis as evaluated by histological examination and inflammatory cytokine profiling (Figure S3).

**Figure 2.**
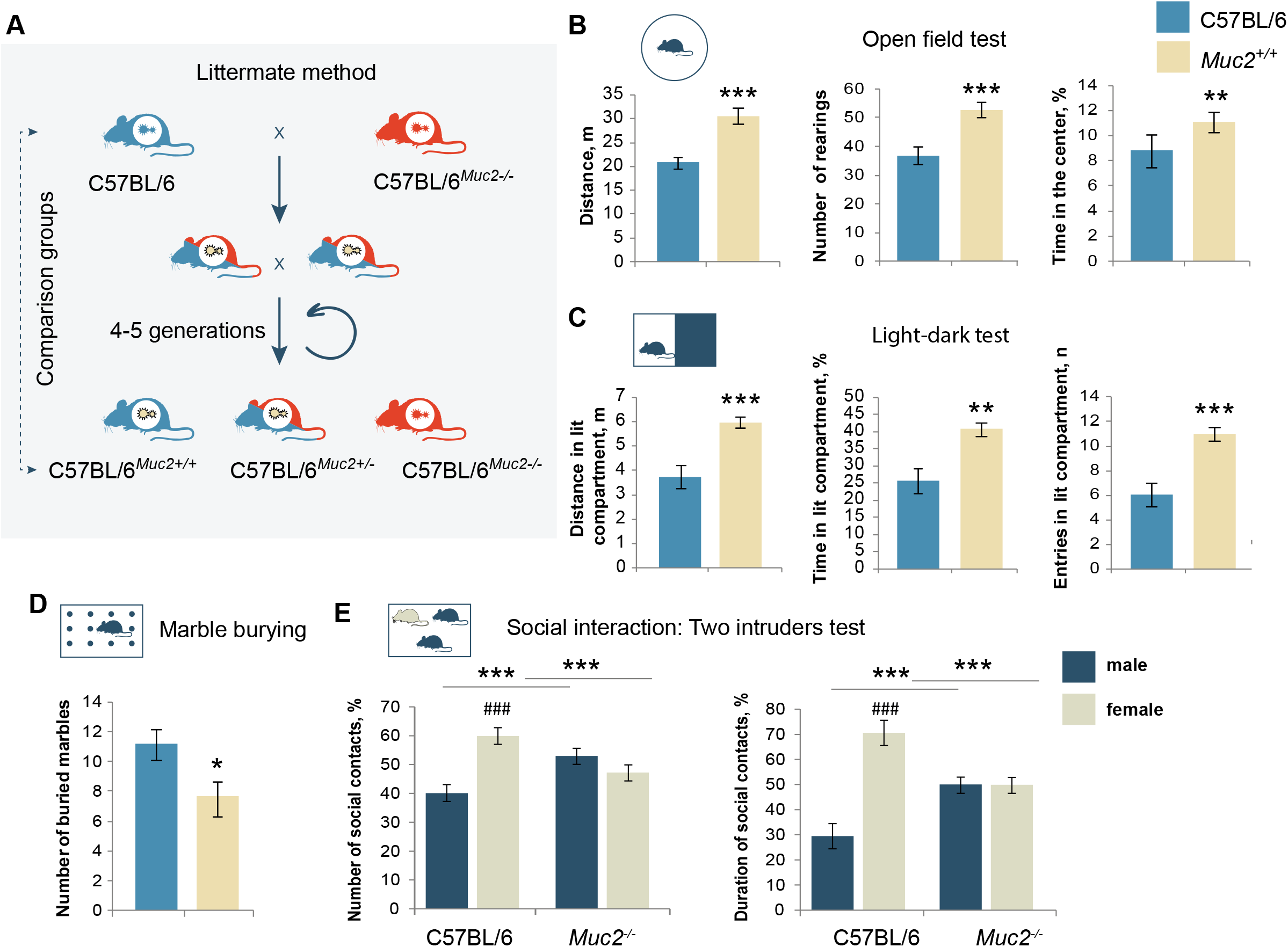
Behavioral traits are associated with microbiota of *Muc2^−/−^* mice. **(A)** The scheme of the littermate co-housing method. **(B)** Open field test (*n* = 18-20). Distance: *t* = −3.895, *p* < 0.001; rearings: *t* = −4.97, *p* < 0.001, Student’s *t*-test; time in the center: Z = −2.997, *p* < 0.001, Mann-Whitney *u* test. (**C)** Light-dark test (*n* = 17-20). Distance: *t* = −4.019, *p* < 0.001; time: *t* = −3.584, *p* < 0.001; entries: *t* = −4.26, *p* < 0.001, Student’s *t*-test. **(D**) Marble burying test (*n* = 18-20). Number of buried marbles: Z = 2.22, *p* = 0.026; Mann-Whitney *u* test. **(E**) Z = 2.22, *p* = 0.026; Mann-Whitney *u* test. **(F)** Two intruders test (*n* = 10-11). Two-way ANOVA revealed a statistically significant interaction between the resident group and the intruder gender (number of contacts: F(1, 38) = 15.490, *p* < 0.001; duration: F(1, 38) = 28.306, *p* < 0.001 Number of contacts (C57BL/6, male *vs*. female): *p*<0.001, Fisher’s LSD test; duration (C57BL/6, male *vs*. female): *p* < 0.001, Fisher’s LSD test. Number of contacts with a male/female intruder *vs*. C57BL/6: *p* = 0.008, Fisher’s LSD test; duration of contact with a male/female intruder *vs*. C57BL/6: *p* < 0.001, Fisher’s LSD test. * = *p* < 0.05, ** = *p* < 0.01, *** = *p* < 0.001, *vs*. C57BL/6. ### = *p* < 0.001, male *vs*. female.

Interestingly, *Muc2^+/+^* mice reconstituted many behavioral features of *Muc2*^−/−^ animals in the open field, light-dark, and marble burying tests (Figure 2). In the open field test, *Muc2^+/+^* mice showed significantly greater activity in comparison with C57BL/6, and reduced anxiety (Figure 2B). In the light-dark test, *Muc2^+/+^* animals were more active in the light compartment as compared to C57BL/6 (Figure 2C). Elevated plus-maze revealed no differences between the test groups (Figure S3). Startle reflex and habituation were on average reduced in *Muc2^+/+^* animals, but these differences were not statistically significant (Figure S3). At the same time, in the marble burying test, *Muc2^+/+^* animals buried fewer marbles consistent with the decreased repetitive behavior and reduced anxiety in mutants (Figure 2D). Social behavior in *Muc2^+/+^* animals was also similar to that of the mutants: *Muc2^+/+^* males made no preferences towards females in terms of social contacts, interacted with a male intruder more than C57BL/6 males, attacked females and mated male intruders (Figure 2E, S3).

We then proposed that if the behavioral pattern in *Muc2^+/+^* animals originates from the *Muc2^−/−^* microbiota transfer, then the direct co-housing with the control C57BL/6 mice should minimize phenotypic differences between these groups. Thus, we co-housed genetically identical *Muc2^+/+^* littermate progeny and C57BL/6 animals to equilibrate their microbiota and found no significant differences in the same panel of behavioral tests (Figure S4). These data demonstrate that the behavioral traits in *Muc2*^−/−^ animals, at least in part, are attributed to the microbiota composition rather than intestinal inflammation itself. Probably, microbiota improves the deteriorating effect of colitis on CNS physiology and masks potential behavioral deficits in mutant mice.

### Manipulation of intestinal microbiota affects behavioral traits associated with Muc2 mutation

We further characterized the intestinal bacterial community in mutant mice along with their wild-type littermates using new generation sequencing of the V3-V4 hypervariable region of *16S rRNA* gene. PCA analysis revealed that mutant animals had substantial differences in the intestinal bacterial community, which were passed via littermate microbiota transfer to the *Muc2^+/+^* mice. Principal component analysis (PCA) revealed significant differences between C57BL/6 and *Muc2^−/−^* and C57BL/6 and *Muc2^+/+^* groups (Figure 3A). Relative abundance of top 10 bacterial genera revealed a decrease in the proportion of *Blautia* and *Escherichia-Schigella*, and an increase in *Akkermansia* upon transfer of *Muc2*-associated microbiota (Figure 3B). Real-time PCR using species-specific primers (Figure S5) confirmed a significant overproliferation of *Akkermansia muciniphila*. To further test the interrelations between behavioral features and microbiota in *Muc2^+/+^* mice, we treated these animals with a broad-spectrum antibiotic amoxicillin combined with a clavulanic acid (AMC) followed by the open field test and the light-dark test. A significant decrease in *A. muciniphila* in the intestine of *Muc2^+/+^* animals was confirmed by real-time PCR (Figure S5). Open field test showed that AMC treatment decreased motor activity and increased anxiety-related behavior (Figure 3C). Light-dark test further confirmed that inhibition of microbiota in *Muc2^+/+^* mice reduced exploratory activity (Figure 3D). These results suggested that inhibition of the intestinal bacteria lead to the normalization of some behavioral traits associated with *Muc2* colitis model.

**Figure 3.**
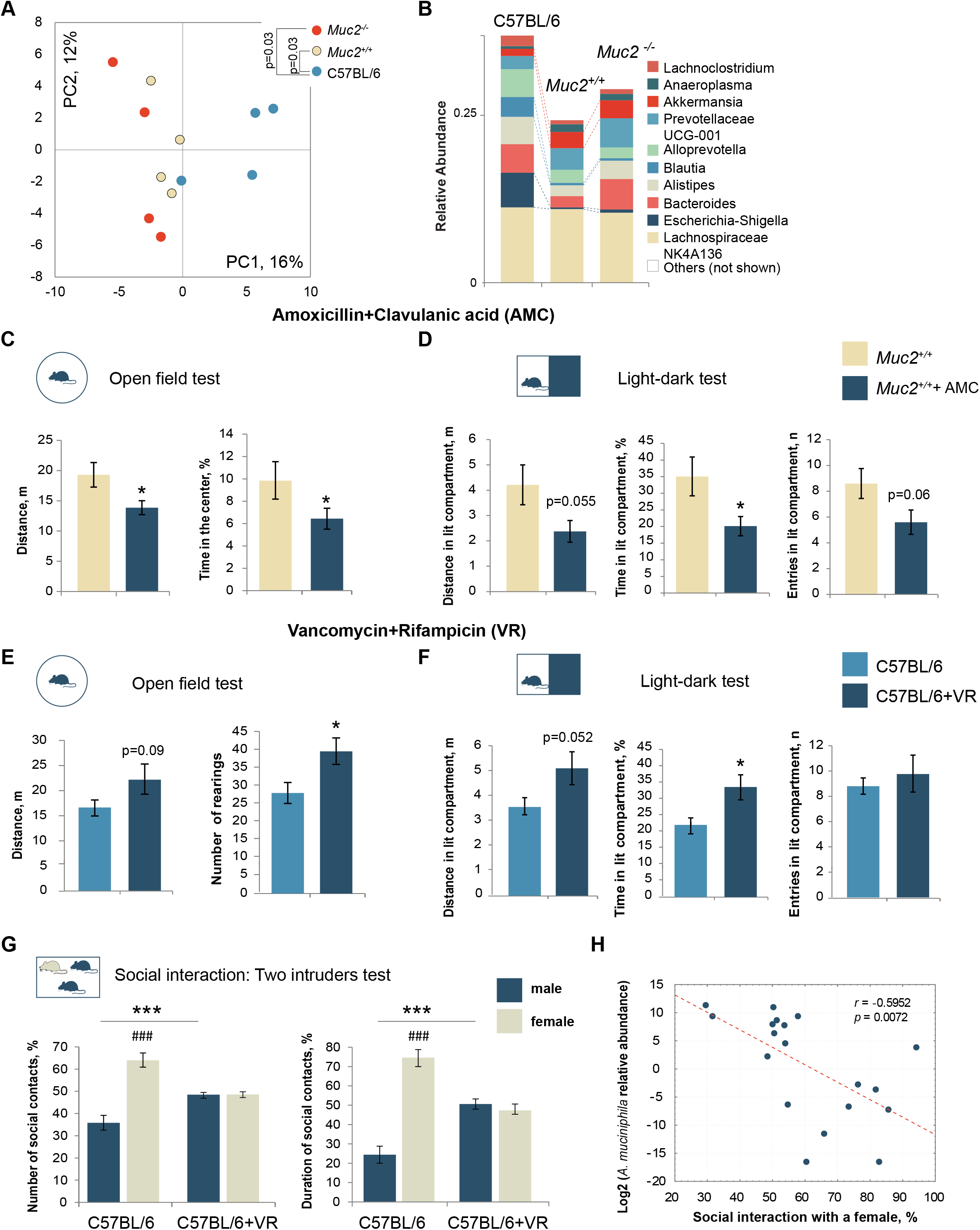
Intestinal microbiota defines behavioral phenotype in the *Muc2* mouse model of colitis. **(A)** PCA of metagenomic data based on top 70 abundant genera (*n* = 4). There was a significant effect of the group in PC1 (*p* = 0.044, Kruskal-Wallis test). There was a significant difference between C57BL/6 and *Muc2^−/−^* and C57BL/6 and *Muc2^+/+^* groups (*Muc2^−/−^*: Z = 2.16, *p* = 0.03; *Muc2^+/+^*: Z = 2.16, *p* = 0.03, Mann–Whitney *u*-test). **(B)** Qualitative changes in average relative abundance of top 10 abundant genera. **(C)** Open field test after AMC treatment (*n* = 10/group). Distance: *t* = 2.34, *p* = 0.03; rearings: *t* = −1.82, *p* = 0.086; time in the center: Z = −1.97, *p* = 0.049; Mann-Whitney *u* test. **(D)** Light-dark test after AMC treatment (*n* = 10/group). In the lit compartment: time: *t* = 2.30, *p* = 0.034; distance: *t* = 2.05, *p* = 0.055; entries: *t* = 2.008, *p* = 0.059, Student’s *t*-test. **(E)** Open field test after VR treatment (*n* = 9/group). Distance: *t* = −1.79, *p* = 0.09; rearings: *t* = −2.47, *p* = 0.025; time in the center: Z = −1.97, *p* = 0.049; Mann-Whitney *u* test. **(F)** Light-dark test after VR treatment (*n* = 9/group). In the lit compartment: time: *t* = −2.553, *p* = 0.022; distance: *t* = −2.09, *p* = 0.052; entries: *t* = −0.643, *p* = 0.529, Student’s *t*-test. **(G)** Two intruders test after VR treatment (*n* = 9-10). Two-way ANOVA revealed a statistically significant interaction between the treatment and the intruder gender (number of contacts: *F*(1, 34) = 28.181, *p*<0.001; duration: F(1, 34) = 56.854, *p*<0.001. Number of contacts (C57BL/6, male *vs*. female): *p* < 0.001, Fisher’s LSD test; duration (C57BL/6, male *vs*. female): *p* < 0.001, Fisher’s LSD test; number of contacts with a male/female intruder *vs*. treatment: *p* < 0.001, Fisher’s LSD test; duration of contact with a male/female intruder *vs*. treatment: *p* = 0.001, Fisher’s LSD test. **(H)** Correlation analysis between social interaction with a female and relative abundance of *A. muciniphila* in the intestinal contents in the control and VR-treated animals: *r* = − 0.595, *p* = 0.007, Pearson’s correlation analysis. * = *p* < 0.05, ** = *p* < 0.01, *** = *p* < 0.001, *vs*. treatment. ### = *p* < 0.001, male *vs*. female.

Next, we reasoned that the behavioral phenotype might be attributed to the abundance of *Akkermansia* as it was the only genus upregulated in *Muc2^−/−^* and *Muc2^+/+^* mice, and *A. muciniphil*a was significantly increased in *Muc2^−/−^* as compared to C57BL/6 animals (Figure 3B, S5). Thus, we aimed to mimic *A. muciniphila* overproliferation in C57BL/6 mice using vancomycin, as it was previously shown to elevate *Akkermansia* (Ray et al., 2020). At the same time, this antibiotic is known for occasional strong outbreaks of *Escherichia coli* proliferation, which might also affect behavior (Jang et al., 2018; Kim et al., 2020; Park et al., 2021). We have shown previously that *E. coli* found in our in-house colony of C57BL/6 is particularly sensitive to rifampicin (Borisova et al., 2020a), so we treated C57BL/6 mice with a combination of vancomycin and rifampicin (VR). PCR analysis revealed that this scheme of antibiotic treatment effectively upregulated *A. muciniphila* in the intestinal contents of C57BL/6 animals and reduced *E.coli* outbreaks (Figure S5). Finally, we performed the open field, light-dark and two intruders tests to analyze the effect of *Akkermansia* expansion on exploratory activity, anxiety-related behavior, and social interactions. In the open field test, RV-treated animals tended to be more active (Figure 3E). The mutant behavioral phenotype was also partially reconstituted in the light-dark test as VR treatment induced more activity in the light compartment (Figure 3F). Even more pronounced was the effect of VR on social behavior: VR-treated males lost preference towards females in the two intruders test, and they interacted with a male intruder more than the control males (Figure 3G). Moreover, the time of interaction with a female negatively correlated with the abundance of *A. muciniphila* in the intestinal contents of the test animals (Figure 3H). Altogether, these results suggest that *A. muciniphila* is a potential regulator of CNS physiology in *Muc2-*associated colitis.

### Muc2-associated microbiota affects blood metabolomic profile and brain glycine level

As microbiota transfer partially reconstitutes behavioral traits in the wild-type littermates in the absence of inflammation, we proposed that metabolomic changes induced by intestinal microbes might mediate the gut-brain interaction in the *Muc2* model of colitis. To test this hypothesis, we performed blood metabolic profiling in *Muc2*^−/−^, *Muc2^+/+^* and C57BL/6 animals using NMR spectroscopy. PCA analysis revealed that *Muc2*^−/−^ littermate co-housing strongly affects host metabolism, and metabolomic profile of the C57BL/6 was significantly different from the two other groups (Figure 4A). Metabolites with similar changes in *Muc2^−/−^* and *Muc2^+/+^* mice are more likely to be involved in regulation of *Muc2*-associated behavioral pattern. Among those were choline, betaine, ketoleucine, 2-hydroxy-isovaleriate, leucine, phenylalanine, glucose, and carnitine (Figure 4B, Table S2).

**Figure 4.**
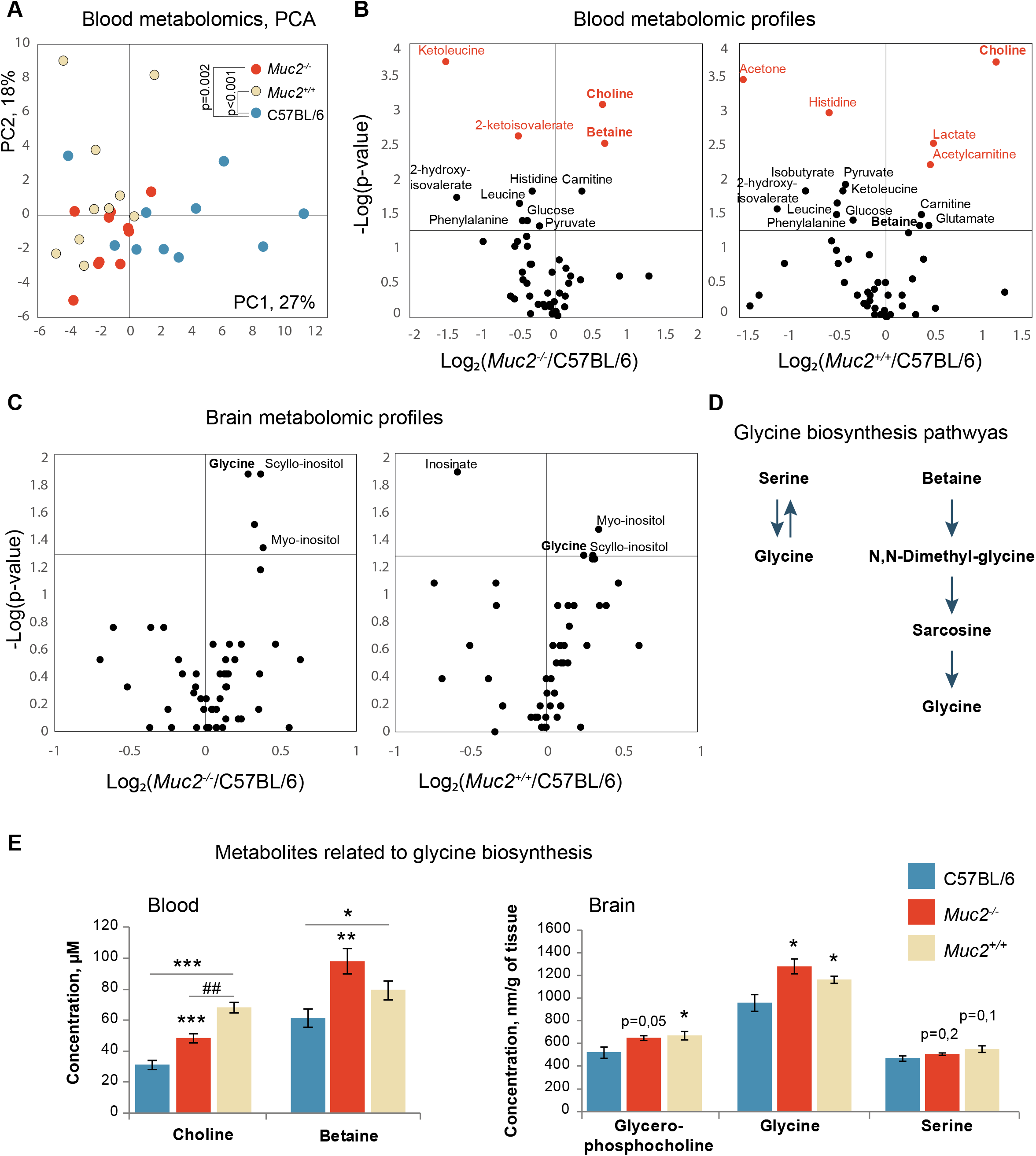
Metabolomic profiling of blood and brain of *Muc2^−/−^* and *Muc2^+/+^* mice. (**A)** PCA analysis of blood NMR metabolic profiles (*n* = 10/group). There was a significant effect of a group on PC1, which accounted for about 26.6% of variance (*F* (2, 27) = 8.84, *p* < 0.001, one-way ANOVA). Metabolomic profile of the C57BL/6 was significantly different from the two other groups (*Muc2*^−/−^: *p* = 0.002; *Muc2^+/+^*: *p* < 0.001, Fisher’s LSD test). (**B)** Volcano plots of blood metabolites as revealed by NMR. Horizontal line depicts a cut-off at *p* = 0.05. Vertical line depicts the ratio of 1. Metabolites with differences at *p* < 0.01 are shown in red. (**C)** Volcano plots of brain metabolites as revealed by NMR (*n* = 5-6). (**D)** The schemes of glycine biosynthesis from serine and betaine. (**E)** The levels of metabolites related to glycine biosynthesis. The data of the Kruskal–Wallis test followed by Mann–Whitney *u*-test are presented in Supplementary tables 2 and 3. * = *p* < 0.05, ** = *p* < 0.01 *vs*. C57BL/6. ## = *p* < 0.01 *vs*. *Muc2^+/+^*.

To further investigate the effect of co-housing on brain metabolism, we performed NMR-based metabolomic profiling of brain tissue from *Muc2*^−/−^, *Muc2^+/+^* and C57BL/6 animals. Based on 50 detected metabolites, PCA did not reveal any overall differences between the groups. However, there were significant changes in some metabolites in *Muc2^−/−^* and *Muc2^+/+^* groups in comparison to C57BL/6 (Figure 4C, Table S3). Inositol (myo- and scyllo- isomers) and glycine were elevated in both test groups (Figure 4C). Glycine was the most interesting among these as it is a neurotransmitter in the CNS and is involved in regulation of anxiety and sensorimotor gating. The major route of glycine biosynthesis is from serine via a reversible reaction catalyzed by Serine hydroxymethyltransferase. Alternatively, glycine can be synthesized from choline via betaine as an intermediate (Figure 4D). Serine was slightly increased in the brain of *Muc2^+/+^* animals, but no significant difference was found for this metabolite (Figure 4E), whereas blood NMR profiles did not reveal a reliable serine signal. At the same time, both blood choline and betaine levels were higher in *Muc2^−/−^* and *Muc2^+/+^* animals (Figure 4E). Brain NMR showed no differences in choline, whereas betaine was not detected (Table S3). However, there was an elevation of glycerophosphocholine in the brain (Figure 4E). Thus, a potential cause of glycine upregulation in the brain might be blood betaine and choline, that are linked to bacterial metabolism (Romano et al., 2017). Given the results of blood and brain NMR, we suggested that glycine might be a potential central mediator of the behavioral phenotype observed in *Muc2* colitis model.

### Pharmacological inhibition of glycine neurotransmission normalizes behavioral changes in Muc2^−/−^ mice

To understand the role of glycine-dependent neurotransmission in regulation of the *Muc2* mouse behavior, we aimed to partially compensate its up-regulation by inhibiting strychnine-sensitive glycine receptors (GlyR) and glycine binding sites in N-methyl-D-aspartate receptors (NMDAR) – the well-known glycine targets. Glycine acts as an inhibitory neurotransmitter via GlyRs, and as a glutamate co-activator at NMDARs. First, we inhibited glycine binding sites at GlyRs via i.p. injection of their inhibitor strychnine to *Muc2^−/−^* male mice. In the open field test, strychnine significantly reduced motor activity (Figure 5A). Likewise, in the light-dark test, there was a decrease in anxiety (Figure 5B). Moreover, strychnine normalized the startle reflex and habituation in mutant males (Figure 5C). Most notably, GlyRs inhibition in *Muc2^−/−^* mice restored social behavior in the two intruders test and reduced overall aggression with no significant effect on the total duration of social contacts (Figure 5D, E).

**Figure 5.**
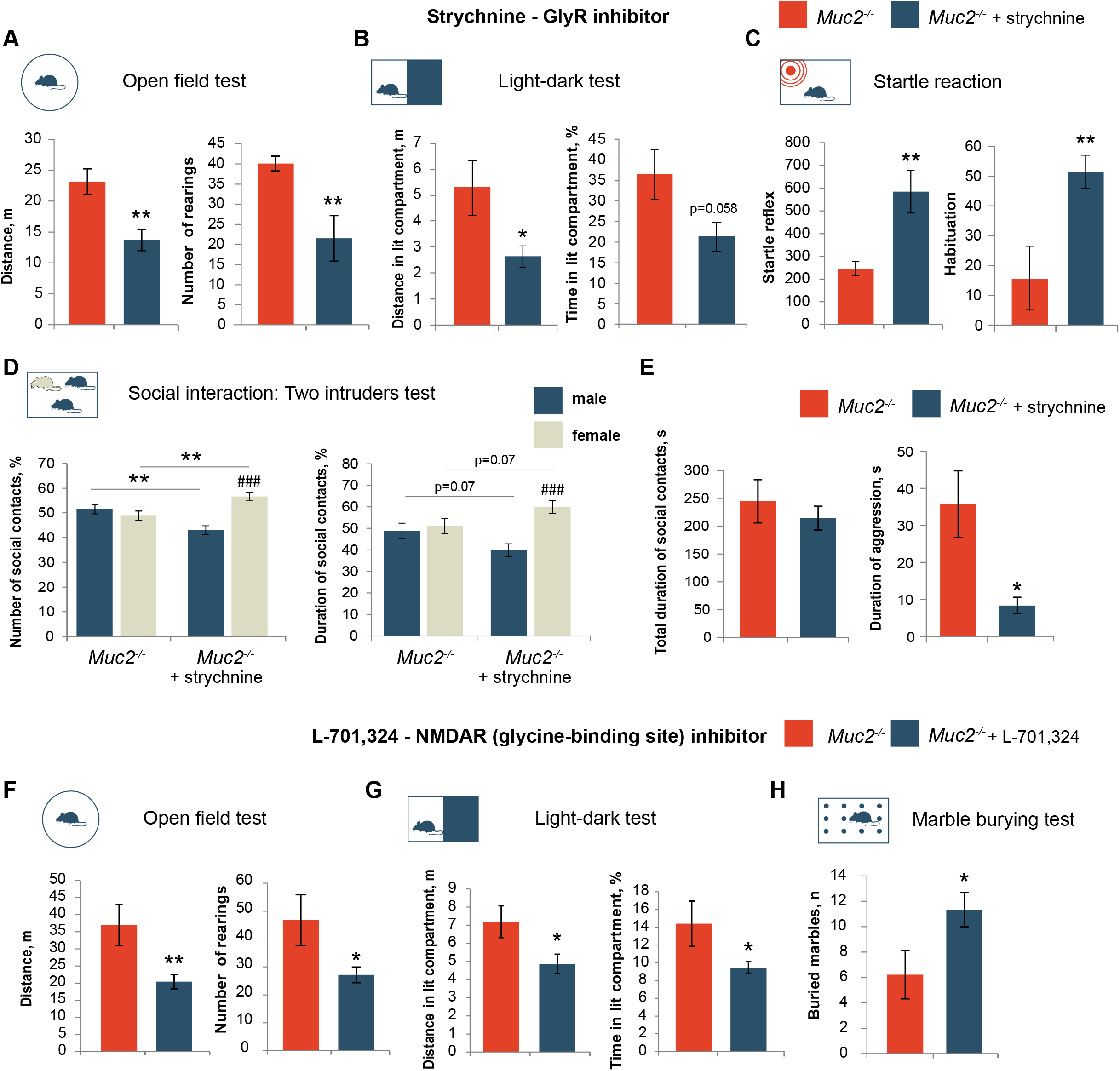
Glycine neurotransmission mediates behavioral abnormalities in *Muc2* model of colitis. **(A)** Open field test after strychnine treatment (*n* = 10-13). Distance: *t* = 3.53, *p* = 0.001; rearings: *t* = 2.897, *p* = 0.009, Student’s *t* -test. **(B)** Light-dark test, (*n* = 7-8). Distance in light compartment: *t* = 2.21, *p* = 0.045; time in light compartment: *t* = 2.08, *p* = 0.058, Student’s *t*-test. **(C)** Startle reflex and habituation (*n* = 10-12; startle: *t* = −3.16, *p* = 0.005; habituation: *t* = −3.09, *p* = 0.006, Student’s *t*-test). **(D)** Two intruders test (*n* = 10/group). Two-way ANOVA revealed a statistically significant interaction between the treatment and the intruder gender (number of contacts: *F*(1, 46)=21.39, *p*<0.001; duration: *F*(1, 46)=6.95, *p* = 0.011. Number of contacts (*Muc2^−/−^* + strychnine, male *vs*. female, *p* < 0.001, Fisher’s LSD test; duration (*Muc2^−/−^* + strychnine, male *vs*. female): *p* < 0.001, Fisher’s LSD test. Number of contacts with a male/female intruder *vs*. treatment, *p* = 0.003, Fisher’s LSD test; duration of contact with a male/female intruder *vs*. treatment, *p* = 0.07, Fisher’s LSD test. **(E)** Two intruders test (*n* = 10/group). Total duration of social contacts (not significant), duration of aggression: *t* = 0.75, *p* = 0.022, Student’s *t*-test. **F**. Open field test after L-701,324 treatment (*n* = 5-11). Distance: *t* = 3.303, *p* = 0.005; rearings: *t* =2.71, *p* = 0.017, Student’s *t* -test. **G**. Light-dark test, (*n* = 5-11). Distance in light compartment: *t* = 2.36, *p* = 0.033; time in light compartment: *t* = −2.56, *p* = 0.023, Student’s *t*-test. **H**. Marble burying test (n = 9/per group, Z = −1.99, *p* = 0.047; Mann-Whitney *u* test). * = *p* < 0.05, *** = *p* < 0.001 *vs*. C57BL/6. ### = *p* < 0.001, male *vs*. female.

To inhibit glycine neurotransmission at NMDA receptors, we used a specific inhibitor of their glycine-binding sites, L-701,324. In the open field test, L-701,324 caused a decrease in activity (Figure 5F). This result was supported by the light-dark test as L-701,324-treated animals spent less time investigating the light compartment (Figure 5G). Interestingly, L-701,324 also normalized marble burying: *Muc2^−/−^* animals buried more marbles upon administration of L-701,324 (Figure 5H). Taken together, both inhibitors ameliorated all behavioral effects related to novelty-induced hyperactivity, anxiety, sensorimotor gating, and social behavior. Therefore, glycine appears to be the major gut-brain axis regulator in the mucus deficiency model of colitis.

## Discussion

*Muc2* knockout mouse model recapitulates key features of chronic intestinal inflammation and demonstrates high clinical and histological score of colitis (Borisova et al., 2020b). At the same time, our behavioral phenotyping revealed no signs of anxiety- or depression-related behavior, as was found previously in other IBD models and human studies (Bercik et al., 2011; Moulton et al., 2019; Neuendorf et al., 2016; Selinger and Bannaga, 2015). This finding suggests that microbiota defines the mutant behavioral profile. Indeed, microbiota transfer via littermate co-housing and combination of antibiotic treatments supported this hypothesis.

Identification of bacterial species associated with mutant behavioral traits revealed an increase in *A. muciniphila*. Interestingly, *A. muciniphila*, the major *Akkermansia* species in the intestine, is a mucin-degrading species and is usually associated with healthy gut and positive effects on the host physiology (McGaughey et al., 2019; Zhai et al., 2019). Its up-regulation in *Muc2* knockout seems discrepant given the nature of this mutation. Probably, the decrease in other commensal bacterial species like *Escherichia* and *Blautia* provide a niche to *Akkermansia* in the gut. At the same time, a number of membrane-bound mucins are present throughout the GI tract (Rodríguez-Piñeiro et al., 2013). It is possible that the lack of Muc2 increases access to other mucins, which might be a preferred mucus source to *Akkermansia*. Consistent with our findings, published data indicate that probiotic treatment with *A. muciniphila* reduces depression, anxiety-related and repetitive behavior and ameliorates motor degeneration, learning and memory deficits in various mouse models (Blacher et al., 2019; Burokas et al., 2017; Ou et al., 2020). At the same time, *Akkermansia* has not been shown to regulate male-male or male-female behaviors, like aggression and courtship. Here we show that intestinal microbiome strongly affects these behaviors, with *A. muciniphila* emerging as a key player in male-male and male-female social interactions.

Metabolic regulation represents one of the major possible communication routs in the gut-brain axis. Our NMR-based metabolomics shows that blood metabolic profiles shist along with *Muc2*-associated microbiome transfer via littermate co-housing, supporting a significant impact of microbiota on host metabolism noticed earlier (Martin et al., 2019; Mithieux, 2018). Brain metabolomic profiles, even though did not show such a drastic association with microbiota, revealed glycine as a potential neurotransmitter responsible for the observed behavior. In our model, inhibition of the glycine binding sites at Gly and NMDA receptors normalized the behavioral profile of *Muc2* knockout animals (Figure 5). Glycine was previously shown to mitigate anxiety-related behaviors, ameliorate obsessive-compulsive disorder, and control sensorimotor gating (Cleveland et al., 2009; File et al., 1987). Most probably, anxiolytic effect of glycine partially masks the effect of inflammation on behavior underlying the lack of depression- and anxiety-like phenotypes in *Muc2* chronic inflammation model. A notable addition to the already known behavioral repertoire of glycine is the involvement of GlyRs in male-male and male-female interactions.

An important question remains on the mechanisms of the central glycine up-regulation and its relation to *A. muciniphila*. Blood metabolomic profiling revealed elevation of choline and betaine in both *Muc2^−/−^* and *Muc2^+/+^* mice. Choline and betaine are precursors of glycine biosynthesis in the glycine-betaine pathway (Razak et al., 2017; Soloway, S.; Stetten, D., 1953). It is possible that elevated levels of these metabolites define high brain glycine. Choline was shown to readily cross the blood-brain barrier (Wurtman et al., 2009) and reduce anxiety-related and repetitive behavior in a mouse model of autism (Agam et al., 2020). Similar effects were shown for the choline downstream metabolic product – betaine (Huang et al., 2019; Kim et al., 2013; Yoshihara et al., 2021). These effects resemble the phenotype observed in *Muc2* model, and therefore elevated choline and betaine might mediate the gut-brain crosstalk. There are a number of potent ways for choline to affect CNS function, including up-regulation of the brain acetylcholine (Cohen and Wurtman, 1975; Koshimura et al., 1990). We have not investigated this neurotransmitter and its effects here as elevated pre-synaptic choline was shown to impair spontaneous alternations in Y-maze and reduce time in open arms in elevated plus-maze (Holmstrand et al., 2014), which is not consistent with our behavioral phenotyping. Moreover, we did not observe choline up-regulation in the brain indicating its rapid peripheral metabolism, which is consistent with the increased blood betaine.

In turn, free choline either comes from the diet or is released by phospholipase D from phosphatidylcholine, known as a main choline depot (Onono and Morris, 2020; Zhaoyu Li 1, 2008). A possible mechanism might involve a shift in the intestinal phospholipid metabolism induced by *Akkermansia* via Toll-like receptor 2 (Plovier et al., 2017). In support of this hypothesis, *Muc2* mutant mice were shown to have deregulated energy pathways and activated lipid metabolism (Tadesse et al., 2017; Ye et al., 2021). So far, it is unclear whether metabolic features of mucin-deficient animals shape their microbial community so that microbiota further promote these changes in a positive feedback manner, or the lack Muc2 predisposes to the microbial changes first. It would be important to investigate this line of relations further, as it might give a possible target in microbiota-induced metabolism to combat neurological consequences of colitis.

## Acknowledgements

This study was supported by the Russian Science Foundation (RSF) Grant #20-74-10022.

## Author Contributions

Formal analysis, M.V.M. and E.N.K.; Funding acquisition, E.N.K.; Investigation, M.V.M., M.A.B., O.A.S., E.A.L., K.M.A., and E.N.K., Methodology, M.V.M., O.A.S., and E.N.K.; Supervision, E.N.K.; Writing – original draft, M.V.M., M.A.B., and E.N.K.; Writing – review & editing, M.V.M., M.A.B., O.A.S., E.A.L., K.M.A., Y.P.T, and E.N.K. All authors have read and agreed to the published version of the manuscript.

## Materials and methods

### Lead contact

Further information and requests for laboratory resources and reagents should be directed to and will be fulfilled by the corresponding author, Elena N. Kozhevnikova

### Materials availability

This study did not generate new unique reagents.

### Data and code availability

All data reported in this paper will be shared by the lead contact upon request.

This paper does not report original code.

## Experimental model and subject details

### Animals

The experiments were performed in the Center for Genetic Resources of Laboratory Animals at the Federal research center Institute of Cytology and Genetics of The Siberian Branch of the Russian Academy of Sciences (ICG SB RAS) and in the Scientific Research Institute of Neurosciences and Medicine (SRINM). All procedures were conducted under Russian legislation according to the standards of Good Laboratory Practice (directive # 267 from 19.06.2003 of the Ministry of Health of the Russian Federation), institutional bioethical committee guidelines and the European Convention for the protection of vertebrate animals. All procedures were approved by the Bioethical committee at SRINM, protocol #8 dated 15.08.2019. All animals had SPF status, which was tested quarterly according to Federation of European laboratory animal science association’s (FELASA) recommendations (Mähler et al., 2014).

The study was conducted using C57BL/6JNskrc (our in-house C57BL/6J sub-colony) and *Muc2^−/−^* mouse strains. C57BL/6JNskrc were recovered from C57BL/6J F1 frozen embryos using SPF CD1 females as foster mothers. *Muc2^−/−^* mice were obtained by the rederivation of previously published *Muc2^tm1Avel^/Muc2^tm1Avel^* mice (Velcich et al., 2002b) with C57BL/6 genetic background (Van der Sluis et al., 2006) in SPF CD1 female mice and back-crossed to C57BL/6JNskrc. Mutant animals and their wild-type littermates (*Muc2^+/+^* mice) were generated by crossing *Muc2^+/−^* females to *Muc2^+/−^* males. In behavioral phenotyping, C57BL/6JNskrc (C57BL/6 further on) mice were used as a control group (Figures 1 and 2). Wild type littermates (*Muc2^+/+^* further on) served as a comparison group where indicated. Ten adult male BALB/c mice were used as a live object in the social preference test, 10 adult male and 10 adult estrus females were used in social tests. The female’s fur on the back was stained with blue food coloring (Cadillac, 2006) to discriminate between males and females in the two intruder test.

Adult 8–14 week-old male mice were housed in groups of the same-sex siblings in individually ventilated cages in 12 h/12 h light/dark photoperiod under standard conditions. Food and water were provided ad libitum. Some SPF *Muc2* mice develop severe intestinal prolapses and exhibit substantial weight loss after weaning. None of these animals was included into the test groups. Sucrose preference test was performed using 8-14 week-old female mice housed under the same conditions.

## Method details

### Co-housing method

Littermate co-housing: *Muc2^−/−^* mice were crossed to C57Bl/6 to generate heterozygous mutants (*Muc2^+/−^*). *Muc2^+/−^* animals were crossed for five generations to obtain *Muc2^+/+^* and *Muc2^−/−^* littermates used in this study.

Direct co-housing was performed in order to control for potential mutations that could accumulate in *Muc2^+/+^* mice during the five generation of *Muc2^+/−^* crosses. It was performed as follows: *Muc2^+/−^* female mice were crossed to *Muc2^+/−^* males. C57BL/6 females were crossed to C57BL/6 males. Pregnant *Muc2^+/−^* and C57BL/6 were housed in separate open cages next to each other with an exchange of dirty litter twice a week. Upon weaning, *Muc2^+/+^* and C57BL/6 progeny were labelled, mixed and co-housed in same cages until behavioral tests were completed (Ridaura et al., 2013).

### Behavioral tests

The order of the behavioral tests with males were as follows: home cage activity, open field, light/dark box, elevated plus maze, marble burying test, Barnes maze, social preference, T-maze, rotarod, tail suspension, forced swim test, startle reflex, active avoidance. For each animal, the tests were performed 3 days apart. Sucrose preference was performed on a separate group of females used for this test only.

All behavioral tests were conducted in the dark time. Three days before the experiment, mice were placed into individual cages to avoid the group effects. For each experiment, a test group and a control group were tested on the same day. In some cases, all groups were divided so that at least some test and control animals are tested on the same day in consecutive days, and the results for each group were combined. The mouse behavior during testing was recorded and processed using Ethovision XT 10 software except for a number of behavioral test that were analyzed manually as specified further.

Animals were euthanized using CO_2_ inhalation. Descending colon and brain samples were taken for nuclear magnetic resonance (NMR) metabolomic profiling and gene expression analyses. Intestinal contents were collected for metagenomic and PCR analyses.

### Home-cage behavior

Home cage behavior was studied using PhenoMaster automated home cage phenotyping setup according to the manufacturer’s instructions. Mice were individually housed for three days in PhenoMaster cages. Locomotion, food and water intake and sleep duration were recorded during the second and the third day of housing. A mouse was considered asleep if no movements were detected in 40 sec or longer (Pack et al., 2007). Animal weight was measured manually.

### Open field test

A mouse was placed in a center of a square plastic setting sized 40 cm x 40 cm with transparent walls and an opaque bottom under white light. Central square 20 cm x 20 cm in size were considered as the center. Animal’s movements were recorded by an overhead video camera and the rearings were detected by a side-view video camera for 6 minutes. Total distance, time spent in the center of the arena and the number of rearings were analyzed using the Ethovision XT10 software.

### Light-dark test

The light-dark test apparatus consisted of a box (42 x 21 x 25 cm) divided into a smaller (one-third) dark compartment and a large (two-thirds) illuminated compartment. Two chambers were connected via a 3 cm x 4cm opening. A mouse was placed in the middle of the dark chamber, facing away from the opening. During 5 min of test, the distance, time and entries into the lit compartment as well as latency to escape after the initial placement were recorded using Ethovision XT10 software (File et al., 2004; Takao and Miyakawa, 2006).

### Elevated plus maze

A mouse was placed in the center of the maze facing an open arm. The distance in open arms, the number of head dips and the time spent in the open arms were measured during the 5-min observation period using Ethovision XT 10 software (Komada et al., 2008).

### Marble burying

Twenty glass marbles (d = 1.0 cm) were evenly distributed on top of a 4 cm layer of sawdust in the plastic animal cage (37 cm x 21 cm x 15 cm). A mouse was placed in the cage and allowed to move freely. After 30 min of exposure, the animal was removed. The number of marbles buried at least two-thirds deep were counted.

### Tail suspension test

The mouse was hung for 6 minutes by the tail with a strip of masking tape on a horizontal bar at a height of 30 cm. The mouse began to move, trying to free itself and attempted to climb on the tail or hung motionless. The total immobility time in seconds was recorded with the Ethovision XT 10 software. The number of animals that successfully climbed on the tail was recorded manually.

### Forced swim test

Animals were placed into a glass cylinder (d=10 cm, h=44 cm) filled up to 25 cm with tap water at a temperature of 25̊C. After 2 min of adaptation, the total time of immobility (sec) was recorded with the Ethovision XT 10 software for 4 min (Yankelevitch-Yahav et al., 2015). An experimenter constantly monitored the animals during the test.

### Rotarod test

The Rotarod device consisted of a metal frame with a motorized rotating assembly of rods. The mouse was placed on the rod facing the direction of rotation followed by the start of rotation. The rotational speed was increased incrementally by 3 rpm every 10 sec. Rotational speed ranged from 0 to a maximum speed of 30 rpm for a maximum period of 2 min. Mice were tested 2 times using the same protocol with an interval of 20 min. The duration of walking before falling and maximal rotational speed before falling were recorded using Rotarod computer software and averaged between the two tests.

### T-maze spontaneous alternation test

T-maze was carried out in red light (28 lux) in a T-shaped apparatus with two closed side compartments (right and left arms) and a start arm separated by automatic doors. A mouse was initially placed in the start arm for 5 sec, after which the start arm door and one goal arm door were opened allowing the animal to examine only one goal arm. The opening goal arm was random for all mice. The door closed upon entering the goal arm, and after 5 seconds, the mouse was returned to its home cage. After that, the apparatus was cleaned to eliminate animal odor traces, and the mouse was placed to the start arm again for 5 sec. Then the start arm door and both goal arm doors were opened so that the animal could examine any goal arm. Once it was selected, the opposite goal arm door closed. After 5 seconds the mouse was returned to its home cage, the apparatus was cleaned, and the latter session was repeated 14 times for each test animal. Maximal duration of a session was 90 seconds. If no choice was made, the animal was returned to its home cage (Deacon and Rawlins, 2006). The total alternation rate was expressed as a percentage of the correct arm selection (opposite to the previous choice) of the total number of sessions with free arm choice.

### Barnes maze

The mouse was placed in a open circular 120 cm-wide platform located 90 cm above the floor with 40 circular holes (8 cm in diameter) evenly pierced around the periphery. The escape box was located under one of the holes, one of four preset locations was randomly chosen for each mouse and did not change during the experiment. The test was carried out in bright white light (about 1000 lux) to stimulate the desire to escape. Distant cues were placed on the walls and floor visible from the platform to help animals navigate the process of searching for the escape hole. The test was composed of four stages as follows:

*Habituation* (1 day, 2 trials of 5 minutes): the mouse was placed close to the target hole, and if after 5 min the animal did not find the escape box, it was carefully guided and left in the escape box for 2 min.

*Training* (4 consecutive days, 2 trials per day for 5 minutes): the mouse was placed in the center of the platform, and the latent time to find the escape box and the distance traveled to the target hole were recorded using Ethovision XT10 software. In case a mouse failed to find a target hole, it was guided there and left in the escape box for 2 minutes. The trial was repeated after 30 minutes.

*Testing* (1 day, 1 trial of 5 min): the mouse was placed in the center of the platform and the latency to escape and the distance traveled to the target hole were recorded using Ethovision XT10 software (Patil et al., 2009; Rosenfeld and Ferguson, 2014; Sunyer et al., 2007).

*Long-term memory retention* was evaluated on day 12 (1 day, 1 trial of 5 min): the mouse was placed in the center of the platform and the latency to escape and the distance traveled to the target hole were recorded using Ethovision XT10 software. No training was conducted between the 5th and 12th day of the experiment (Patil et al., 2009).

### Sucrose preference test

Sucrose preference was evaluated in the IntelliCage setting. The IntelliCage apparatus consists of a polycarbonate cage (58 cm x 40 x 20.5 cm) with two drinking bottles in each corner and equipped with automatic doors and sensors recording entries of an animal, nose pokes, and licking. The cage floor was covered with the sawdust and used as home cage starting the 4^th^ day of the experiment. The female mice were used in order to avoid aggressive behavior after placement of non-sibling adult animals into the setting. Testing was carried out in two apparatus cages: the first cage contained C57BL/6 females, and the second cage - *Muc2^−/−^* females. Each mouse was labelled with a code of an implanted radio-frequency identification chip.

The following protocol was used:

*Chip adaptation* – 3 days after chip implantation female mice we kept in their home cages.

*Adaptation to IntelliCage* - 7 days. Mice were placed in the IntelliCage supplied with drinking water and food ad libitum. All doors in corners were open and animals had free access to water.

*Sucrose preference* - 5 days. Drinking water was replaced with 2% sucrose in 2 corners out of 4, all doors remained open (Dere et al., 2018; Krackow et al., 2010). The IntelliCage Software automatically recorded entries and licks. Sucrose preference was calculated as the number of sucrose licks in percent of the total number of licks.

### Active Avoidance test

The Gemini Avoidance System apparatus consisted of two automated shuttle-boxes, each one was divided into two 20 x 10 cm compartments, connected with a 3 x 3 cm opening. At the beginning of each test day, an animal was given a 5-minute adaptation period in the apparatus and was allowed to move freely. The location of the animal was detected by sensors and recorded automatically.

Light (10 W) was used as a conditional stimulus 5 seconds prior an unconditioned stimulus - a foot shock (0,4 mA, 5 sec). The intertrial interval was 30 sec. Training consisted of 5 days, each day a 50-trial avoidance sessions were performed. Each trial was classified as an avoidance response (crossing to the other side before the onset of the shock), an escape response (crossing to the other side after the onset of the shock), or a null response (remaining in the original compartment and receiving 5 sec of shock) (Sparkman et al., 2005). The results are presented as an average number of avoidance reactions per day per group.

### Startle reflex, prepulse inhibition (PPI), habituation

The startle reflex, PPI and habituation were evaluated using an SR-Pilot device consisting of a plastic chamber with build-in piezoelectric sensors for movement detection and a controlled source of sound. The session consisted of 5 blocks of 64 trials in total as follows:

*Block 1, acclimation trials 1-6*: 5 min acclimation at background noise level (65 dB);

*Block 2, trials 1-6*: six pulse-alone trials (120 dB, 40 msec);

*Block 3, trials 7-32*: 26 pulse-alone trial (120dB), prepulse (either 69, 73 or 81 dB) + pulse alone trial (120 dB), and NOSTIM trials in a pseudorandom order;

*Block 4, trials 33-58*: 26 pulse-alone trial (120dB), prepulse (either 69, 73 or 81 dB) + pulse alone trial (120 dB), and NOSTIM trials in a pseudorandom order;

*Block 5, trials 1-6*: six pulse-alone trials (120 dB, 40 msec).

PP73 represented prepulse (73 dB, 20 msec) + pulse-alone trials (120 dB, 40 msec), PP81 represented prepulse (81 dB, 20 msec) + pulse-alone trials (120 dB, 40 msec), and NOSTIM trials represented record-only trials. The prepulses were presented 100 msec before the onset of pulse-alone stimuli. The inter-trial intervals lasted for about 15 sec.

The average value for pulse alone in the blocks 3 and 4 were used to calculate Startle reflex. The difference between average values for pulse alone in blocks 2 and 5 were used to calculate Habituation. The difference between average values for pulse alone and prepulse+pulse in the blocks 3 and 4 were used to calculate PPI (Geyer and Dulawa, 2003):

Startle reflex = average (pulse-alone in blocks №3 and 4), expressed in arbitrary units;

PPI = 100 × [pulse-alone(block №3,4) – (prepulse + pulse)]/pulse-alone(blocks №3 and 4), expressed in %;

Habituation = 100× [pulse-alone(block №2) – pulse-alone(block №5)]/pulse-alone (block №2), expressed in %.

### Social odor preference test (SOPT)

The social preference test was conducted according to the protocol described by Malkova with co-authors (Malkova et al., 2012) under red light in a rectangular test arena (60 cm x 40 cm) separated into three equal chambers (20 cm x 40 cm, left, central and right) with a 6 cm x 6 cm openings. An animal was placed in the central chamber and was allowed to move within the arena for 5 minutes for acclimation. An unfamiliar BALB/c male mouse was placed in one side chamber under a wire cup, whereas a white cotton ball of comparable size was placed under similar cup into the opposite chamber. Time spent in each chamber was analyzed automatically using Ethovision XT10 software. Social preference coefficient was calculated as (time spent in the “animal” chamber) / (time spent in the “cotton” chamber). Choosing a chamber to place a BALB/c mouse was random.

### Resident-intruder tests

Test males were single-housed for at least 3 days prior to the test. Three month-old sexually experienced BALB/c animals were used as intruders. Either a female, a male or both intruders were placed in the test animal’s homecage for 15 minutes (Liu et al., 2011; Zolotykh and Kozhevnikova, 2017), and social interactions were recorded by a video camera. Attacks and matings were counted manually during the test by two independent observes and averaged out. Duration and quantity of sniffing and licking was scored manually using video recordings by the two independent observes and averaged out. The female intruder test was performed at first followed by the male intruder test, and the two intruders test was performed last. The inter-test time was at least 2 days for each animal.

### Metagenomic analysis

For the gut metagenomic analysis, intestinal contents (the content of small intestine, cecum, colon and a fecal sample) were collected in Inhibitex buffer, and total DNA was extracted using QIAamp DNA Stool Mini Kit according to manufacturer’s instructions (*n*(*C57BL/6*) = 4, *n*(*Muc2^+/+^*) = 4, *n*(*Muc2^−/−^*) = 4). Metagenomic analysis was performed by a commercially available service at Novogene (https://en.novogene.com) as follows. The *16S rRNA* V3-V4 region was used for barcoded amplification using Phusion High-Fidelity PCR Master Mix, and PCR-products were mixed in equal ratios. The mixture was purified with Qiagen Gel Extraction Kit. The libraries were generated with NEBNext UltraTM DNA Library Prep Kit for Illumina sequencing and analyzed by the Illumina platform. Data analyses were performed using Uparse software. Sequences with ≥97% similarity were assigned to the same operational taxonomic units (OTUs). Beta diversity analysis was used to measure differences between samples in terms of species complexity. Beta diversity of both: weighted and unweighted unifrac were calculated by QIIME software (Version 1.7.0). Principal component analysis (PCA) preceded cluster analysis to reduce the dimension of the original variables using the FactoMineR package and ggplot2 package in R software (Version 2.15.3). Differences in taxa abundance between groups were shown for the average top ten taxa in each group using normalized reads for each OTU. PCA analysis was performed using top 70 genera, identified in at least one of the test groups, top two PCs in terms of variance coverage were used for data plotting and identification of significant differences among the groups. The data is currently in the process of deposition to the public database and is available for the reviewers at https://data.mendeley.com/datasets/y85tx8hzdh/draft?a=5fde3ad3-1078-4911-9523-92a361805999.

### Metabolite extraction and nuclear magnetic resonance (NMR) spectroscopy

The metabolomic composition of the sample was analyzed at the Center of Collective Use «Mass spectrometric investigations» SB RAS (Novosibirsk, Russia) by high-resolution ^1^H NMR spectroscopy. The extraction of metabolites from serum was performed by using a short sample preparation protocol earlier evaluated for quantitative NMR-based metabolomics (Borisova et al., 2020a; Snytnikova et al., 2019). Namely, 100 μL of ice-cold methanol and 100 μL of ice-cold chloroform were added to 100 μL of serum and vortexed for 30 s, kept on ice for 10 min, and incubated at −20 °C for 30 min. The mixtures were centrifuged at 12,000 rpm and at 4 °C for 30 min to pellet proteins. The top hydrophilic fraction was collected to fresh vials and lyophilized using vacuum concentrator.

To obtain protein-free extract of metabolites from the mouse brain, we used the following sample preparation protocol earlier evaluated for quantitative NMR-based metabolomics (Glinskikh et al., 2021). Briefly, frozen brain tissue was weighed and homogenized using a glass homogenizer in cold (−20 °C) water/methanol/chloroform mixture in a ratio 1:2:2 (v/v; 1600 μL of solvent mixture per 150 mg of wet tissue), vortexed for 30 seconds, kept on ice for 10 minutes, and incubated at −20 °C for 20 minutes. The mixtures were centrifuged at 12000 rpm, 4 °C for 30 minutes to pellet proteins. The top hydrophilic fraction was collected to fresh vials and lyophilized using vacuum concentrator.

Dried extracts were re-dissolved in 600 μL of D2O containing 6×10-6 M DSS as an internal standard and 20 mM deuterated phosphate buffer to maintain pH 7.4. The ^1^H NMR measurements were carried out with the use of the AVANCE III HD 700 MHz NMR spectrometer equipped with a 16.44 T Ascend cryomagnet. The proton NMR spectra for each sample were obtained with 64 accumulations. Temperature of the sample during the data acquisition was kept at 25 °C, the detection pulse was 90 degree. The repetition time between scans was 25 s to allow for the relaxation of all spins. Low power radiation at the water resonance frequency was applied prior to acquisition to presaturate the water signal. The pulse sequence zgpr was applied.

The collected NMR spectra were manually phased and baseline corrected. Signal processing and integration was performed using MestReNova V.12 software. We used DSS at a concentration of 6 μM as reference for chemical shift and determination of the metabolite concentration. The metabolite resonance assignments were made by comparison with data of Human Metabolome Database (Wishart et al., 2018) (http://www.hmdb.ca) and our own experience in the metabolomic profiling (Snytnikova et al., 2017a, 2017b; Yanshole et al., 2014, 2019), or by adding reference compounds whenever needed. The concentration of metabolites in the samples was calculated by the integration of the peak area of a metabolite respectively to DSS added to the sample. Amino acids, peptides, organic acids, alcohols, nucleotides, osmolytes, energy metabolism products were identified. The detailed description of metabolite quantification and validation are presented in our previous work (Glinskikh et al., 2021). The data is currently in the process of deposition to the public database and is available for the reviewers at amdb.online/amdb/experiments/152/ (blood serum profiles), amdb.online/amdb/experiments/154/ (brain profiles).

### Real-time PCR

Fecal DNA was purified from fecal pellets using QIAamp DNA Stool Mini Kit according to the manufacturer’s recommendations. Bacterial abundance was measured by real-time PCR using BioMaster HS-qPCR SYBR Blue, 5 μL of fecal DNA and 300 nM specific primers (Supplementary Table 1). The data was normalized to *16SrRNA* as ΔCt = 2^(Ct_*16SrRNA*_ - Ct_bacterium of interest_) and shown as log_10_(ΔCt) (Borisova et al., 2020a). For samples after antibiotic treatment, the data was normalized to *28SrRNA* as ΔCt = 2^(Ct_*28SrRNA*_ - Ct_bacterium of interest_) and shown as log_10_(ΔCt) or log_2_(ΔCt). For gene expression analyses, total RNA from colonic samples was purified using TRIzol reagent. One μg of RNA was used in reverse transcription reaction, cDNA was synthesized using M-MuLV reverse transcriptase according to the manufacturer’s recommendations. The reaction volume of 20 μL was then diluted up to 100 μL with deionized water and used for PCR. Real-time PCR reaction was prepared using a BioMaster HS-qPCR SYBR Blue, 5 μL of cDNA, and 250 nM specific primers. Gene expression was normalized to *Tubb5* mRNA as ΔCt = 2^(Ct_*Tubb5* mRNA_ − Ct_gene of interest mRNA_).

### Drug treatments

Vancomycin was diluted in drinking water at a concentration of 0.5 g/L. The animals of the experimental group (C57BL/6+VR) received vancomycin solution in drinking water for 2 weeks, rifampicin at a concentration of 0,075 g/L has been added in the last 8 days. During the testing period, animals continued to drink vancomycin + rifampicin solution. Control animals C57BL/6 received regular drinking water. Feces for *A. muciniphila* detection were collected in three consecutive days starting the day of the first behavioral test.

Amoxicillin + clavulanic acid were diluted in drinking water at a concentration of 0.45 mg of amoxicillin and 0.15 mg of clavulanic acid per ml of water. *Muc2^+/+^* (*Muc2^+/+^*+AMC) animals received this solution for 2 weeks and during the testing period. Feces for *A. muciniphila* detection were collected in three consecutive days starting the day of the first behavioral test. *Muc2^−/−^* animals were also used for antibiotic treatment, but were withdrawn from this experiment due to excessive weight loss and high death rate.

Strychnine was administered intraperitoneally (i.p.) to *Muc2^−/−^* animals (*Muc2^−/−^* + strychnine) 15 minutes before the test (Kehne et al., 1981) at a dose of 0.75 mg/kg (the dose was selected empirically according to the published data (Commissaris et al., 1998; Kehne et al., 1981), in a volume of 10 ml/kg. Control animals were injected with the same volume of saline. The interval between the tests was 3 days.

The selective antagonist of the NMDA receptor glycine site L-701,324 was dissolved in 99% saline, 1% DMSO (vehicle) and injected i.p. 45 minutes before the behavioral test in a dose of 10 mg/kg. Control mice received 1% DMSO in a volume of 10 mL/kg. The interval between the tests was 3 days.

### Histological and clinical scores

Paraffin sections (4 μm) were stained with H&E (Hematoxylin and Eosin) stain. The sections were examined in a blinded manner. Hyperplasia was defined as the percentage of cells per crypt above the control. Erosion was defined as the area of epithelium where the crypt structure was lost and shown as the percentage per section. Hyperplasia scoring: 0 - <10%; 1 - 10–50%; 2 - 51–100%; 3 - >100%). PMN cell infiltration scores: 0 - none, 1 - mild, 2 - moderate, 3 - severe). Scores for erosion (percentage of area involved): 0 - <1%, 1 - 1–15%, 2 - 16–30%, 3 - 31–45%, 4 - 46–100%). The maximum score (Total score) that could result from this scoring was 10. For clinical scores, animal’s weight, stool consistency and fecal blood were evaluated manually. Weight loss scoring: 0 – none, 1 – 0-17%, 2 – 18-35%, 3 - >35%. Stool consistency and fecal blood scoring: 0 – normal droppings, 1 – loose droppings, 2 – diarrhea and 0 – no blood, 1 – visible blood in rectum, 2 – visible blood on fur. The total score reflects the sum of all scores.

## Data Analysis and Statistics

### Statistical analysis

The data are presented as mean±stadard error of mean, bacterial PCR is shown as data points. The data were tested for normality using the Kolmogorov-Smirnov test. Normally distributed data were analyzed by Student’s t-test test for independent samples. Data involving multiple factors were analyzed using analysis of variance (ANOVA) followed by Fisher LSD test. Not normally distributed data were processed using Kruskal–Wallis test followed by Mann– Whitney *u*-test. Active avoidance was analyzed using Friedman test for repeated measures. Tail suspension results were analyzed using χ2 test. Principal component analysis (PCA) was performed in STATISTICA12 software, data significance in principal components (PC) was evaluated using ANOVA followed by Fisher LSD test (blood metabolomics) or using Kruskal– Wallis test followed by Mann–Whitney *u*-test (brain metabolomics and metagenomics). Correlation was analyzed by Pearson correlation analysis. The value of *p* < 0.05 was considered significant.

## Supplemental figure legends

**Supplementary figure 1. Depressive-like behavior, motor skills and memory retention in *Muc2^−/−^* mice. Behavioral phenotyping of *Muc2^−/−^* animals. (A)** Elevated plus-maze (*n* = 20/group). Number of peeks down: Z = −3.02, *p* = 0.001, Mann-Whitney *u* test. **(B)** Social preference (*n* = 12 /group). **(C)** Forced swim test (*n* = 10-12). **(D)** Sucrose preference (*n* = 11-12). **(E)** Tail suspension (*n* = 11-14). Number of animals that climbed on the tail: *p* < 0.001, χ2 test**. (F)** Rotarod test (*n* = 17-20). Latency to fall: *t* = −2.766, *p* = 0.009; rotational speed: *t* = −2.716, *p* = 0.01, Student’s *t*-test. **(G)** T-maze (*n* = 8-9). **(H)** Barnes maze (*n* = 10-12). There was the influence of the test day on the distance to the target hole (*F* (2, 74) = 33.66, *p* < 0.001) and on the latent time to the escape box (*F* (2, 74) = 71.39, *p* < 0.001 repeated measures ANOVA). Distance to the escape box as compared to day 1: day 5, *p* < 0.001; day 12, *p* < 0.001, Fisher’s LSD test. Escape box latency as compared to day 1: day 5, *p* < 0.001; day 12, *p* < 0.001, Fisher’s LSD test. There was a significant interaction of day and group factors (*F* (2, 74) = 3.14, *p* = 0.049). Latent time to escape box on day 1 as compared to C57BL/6 (*p* = 0.011, Fisher’s LSD test). **(I)** Active avoidance (*n* = 10-11). Number of active avoidances as compared to day 1: *Muc2^−/−^*: χ2 (11) = 11.23, *p* = 0.037; C57BL/6 (χ2 (10) = 18.2, *p* = 0.001, Friedman test, Fig. 2g). * = *p* < 0.05, ** = *p* < 0.01, *** = *p* < 0.001, *vs*. C57BL/6. ### = *p* < 0.001, *vs*. day 1.

**Supplementary figure 2. Social behavior in *Muc2^−/−^* mice. (A)** Male intruder test (*n* = 15-16). Number of sniffing acts: *t* = −2.323, *p* = 0.027, Student’s *t*-test; number of animals with aggressive behavior: *p* = 0.019, χ2 test; number of mating animals: *p* = 0.078, χ2 test. **(B)** Female intruder test (*n* = 16/group). Number of mating animals s: *p* = 0.012, χ2 test. **(C)** Two intruder test (*n* = 10-11). Total social contacts number: *t* = −2.018, *p* = 0.049, Student’s *t*-test; number of animals mating a male (*n* = 7-10): *p* = 0.01, χ2 test; number of animals attacking a female (*n* = 7-10): *p* = 0.002, χ2 test. * = *p* < 0.05, ** = *p* < 0.01 *vs*. C57BL/6.

**Supplementary figure 3. Gene expression and histological analyses of the descending colon in *Muc2^+/+^* mice revealed no inflammatory response. (A)** PCR analysis of pro-inflammatory genes in *Muc2^+/+^* animals as compared to C57BL/6 (*n* = 7/group). Z = 2.468, *p* = 0.014, Mann-Whitney *u*-test. **(B)** Histological (*n* = 6/group) and clinical (*n* = 8/group) scores in *Muc2^+/+^* animals. Clinical scores for *Muc2^−/−^* animals are shown as a non-zero comparison: Z = −3.308, *p* = 0.001, Mann-Whitney *u* test. **(C)** Representative pictures of the descending colon from C57BL/6, *Muc2^+/+^*, and *Muc2^−/−^* mice. Scale bar 100 = μm. **(D)** Elevated plus-maze (*n* = 9-10). **(E)** Startle reflex (*n* = 18/group). **(F)** Two intruders test (*n* = 10-11). Total social contacts number: *t* = −2.88, *p* = 0.009, Student’s *t*-test; number of animals mating a male (*n* = 8-10): *p* = 0.023, χ2 test; number of animals attacking a female (*n* = 8-10): *p* = 0.000, χ2 test. * = *p* < 0.05, ** = *p* < 0.01, *** = *p* < 0.001, *vs*. C57BL/6.

**Supplementary figure 4. *Muc2^+/+^* littermate behavior after co-housing with C57BL/6 mice. (A)** The scheme of direct co-housing method. **(B)** Open field test (*n* = 10-11). **(C)** Light-dark test (*n* = 10-11). Entries in lit compartment: *t* = 2.481, *p* = 0.023, Student’s *t*-test. **(D)** Elevated plus-maze (*n* = 9-10). **(E)** Marble burying test (*n* = 10-11). **(F)** Startle reflex (n = 9-10). **(G)** Two intruders test (*n* = 9-10). There was a statistically significant effect of the intruder gender (duration of contacts: *F*(1, 34) = 67.856, *p* < 0.001, two-way ANOVA). Duration of contacts (C57BL/6, male *vs*. female): *p* < 0.001, Fisher’s LSD test. Duration of contacts (*Muc2^+/+^*, male *vs*. female): *p* < 0.001, Fisher’s LSD test * = *p* < 0.05 *vs*. C57BL/6. ### = *p* < 0.001, male *vs*. female.

**Supplementary figure 5. PCR analysis of *A. muciniphila* and *E. coli*. (A)** PCR analysis of *A. muciniphila* in the intestinal contents normalized to *16S rRNA* (*n* = 9-10). Kruskal-Wallis test revealed a significant effect of the group on *Akkermansia* (H(2, N= 29) = 10.803, *p* = 0.005, Kruskal-Wallis test). *Muc2^−/−^*: Z = −2.95, *p* = 0.003; *Muc2^+/+^*: Z = −2.944, *p* = 0.003; Mann-Whitney *u* test. **(B)** PCR analysis of *A. muciniphila* in the intestinal contents after treatment with amoxicillin and clavulanic acid (AMC: Z = −3.742, *p* < 0.001 *vs*. *Muc2^+/+^*) and with vancomycin and rifampicin (VR: Z = −3.593, *p* < 0.001 *vs*. C57BL/6) normalized to *28S rRNA*, *n* = 9-10. **(C)** PCR analysis of the intestinal contents normalized to *28S rRNA* (*n* = 9-10). Kruskal-Wallis test revealed a significant effect of the treatment: H(2, N= 29) = 12.317, *p* = 0.002. *E. coli* in the intestinal contents after treatment with either vancomycin (V: Z = −3.062, *p* = 0.002 *vs*. C57BL/6) or with vancomycin and rifampicin (VR: Z = −2.735, *p* =0.006 Mann-Whitney *u* test). ** = *p* < 0.01, *** = *p* < 0.001, *vs*. C57BL/6. ### = *p* < 0.001, *vs*. *Muc2^+/+^*.

