## Supplemental figures 1-5 for "Colitis-associated intestinal microbiota regulates brain glycine and host behavior in mice"

Supplementary figure 1

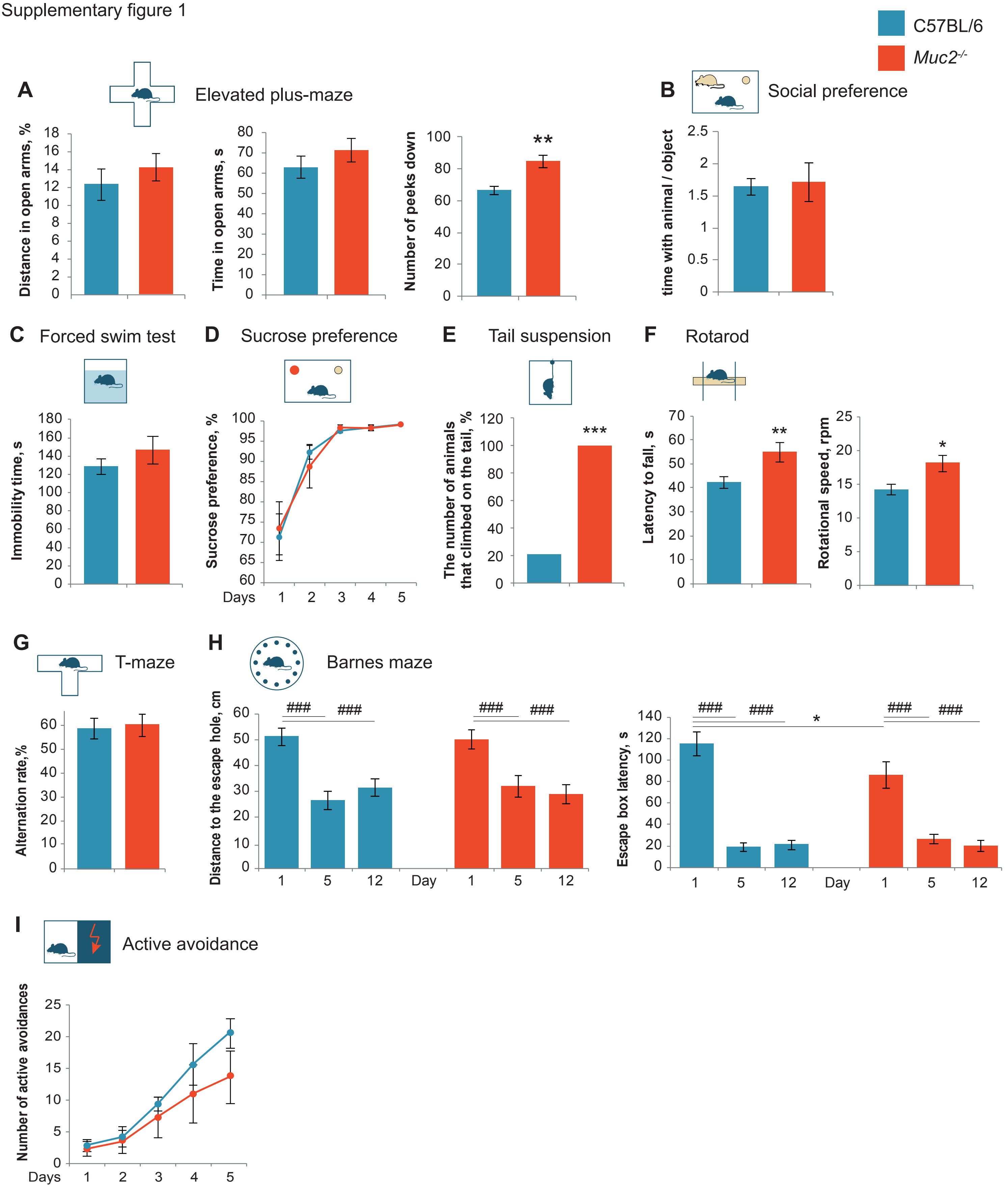

Supplementary figure 2

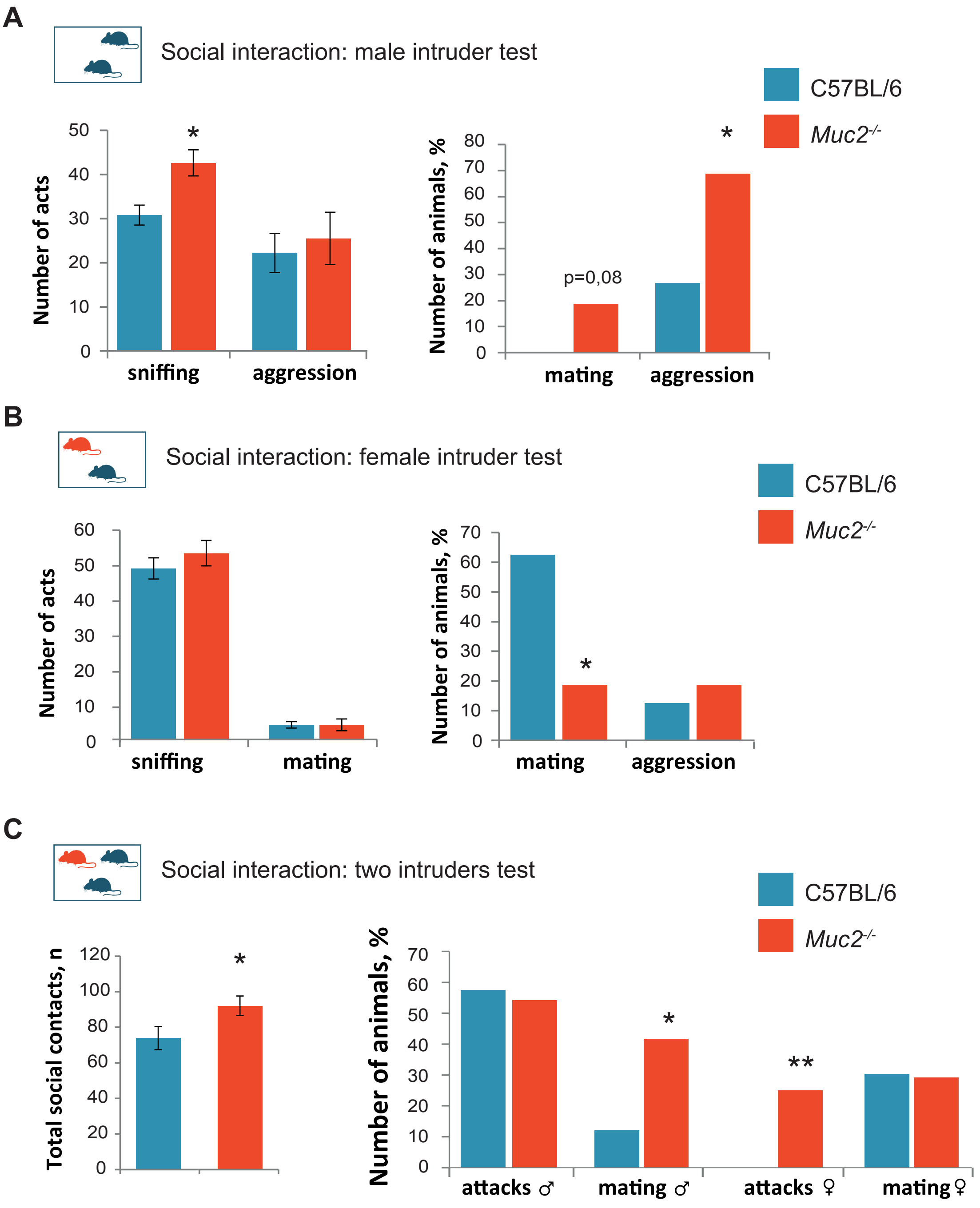

Supplementary figure 3

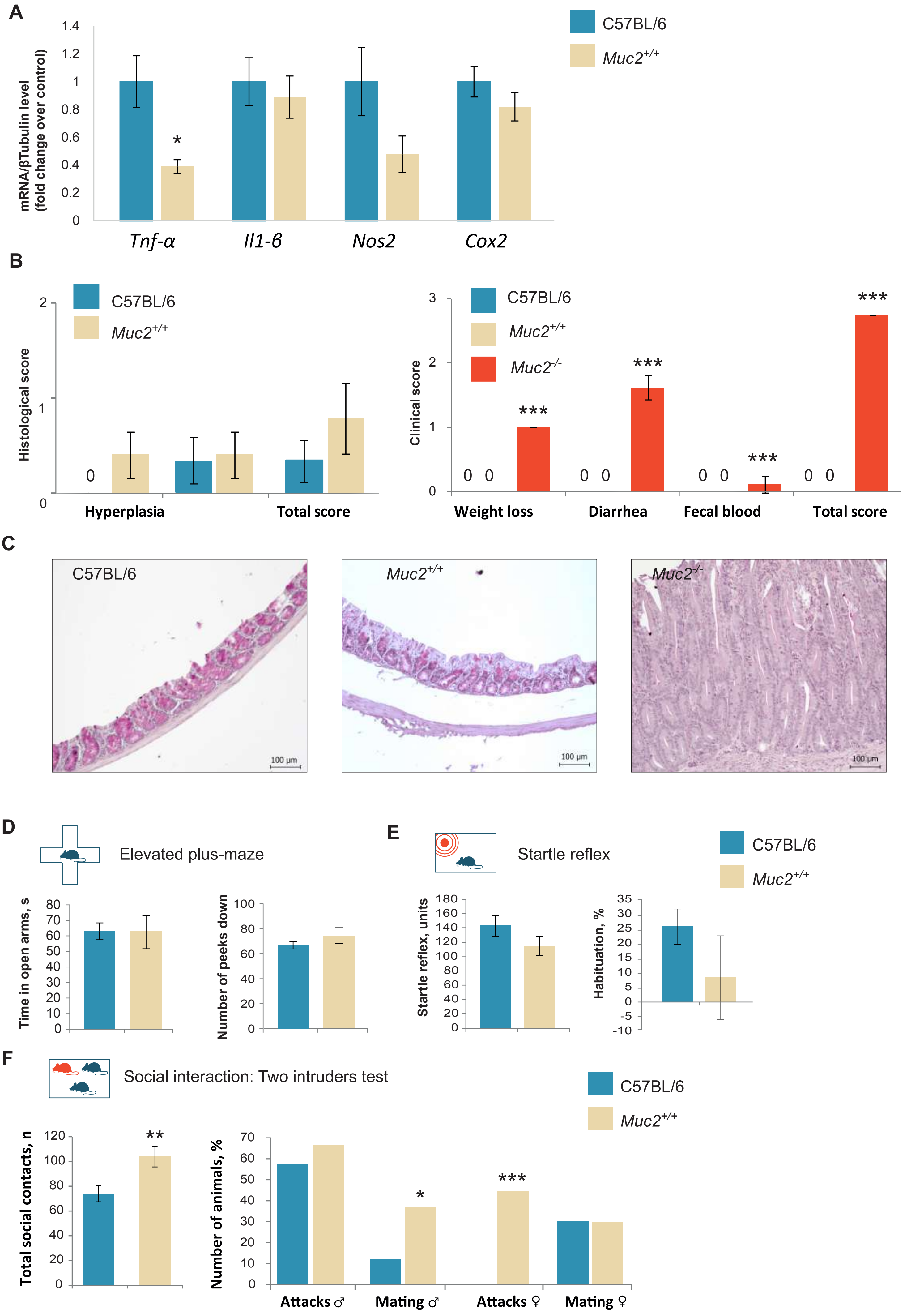

Supplementary figure 4

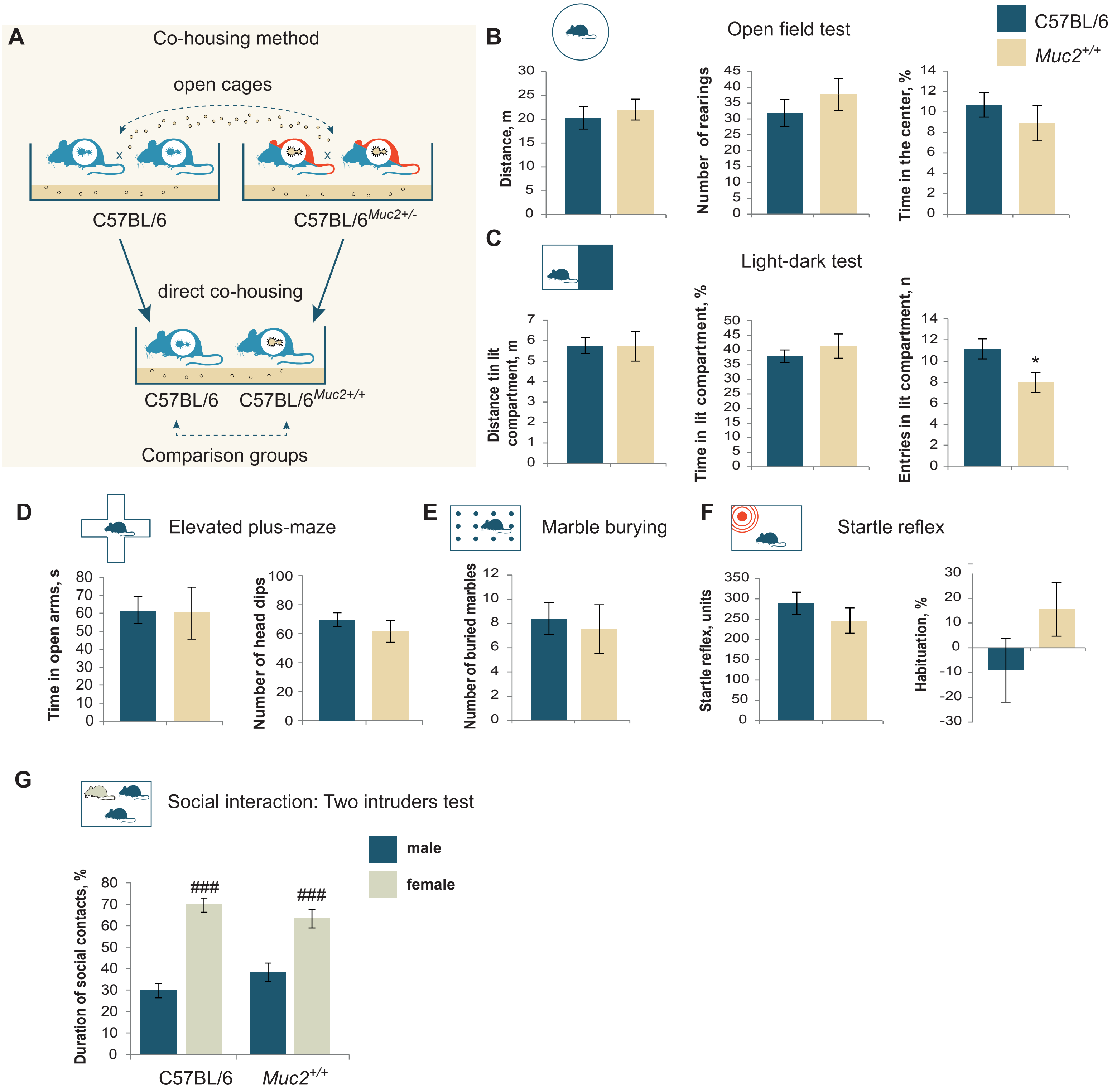

Supplementary figure 5

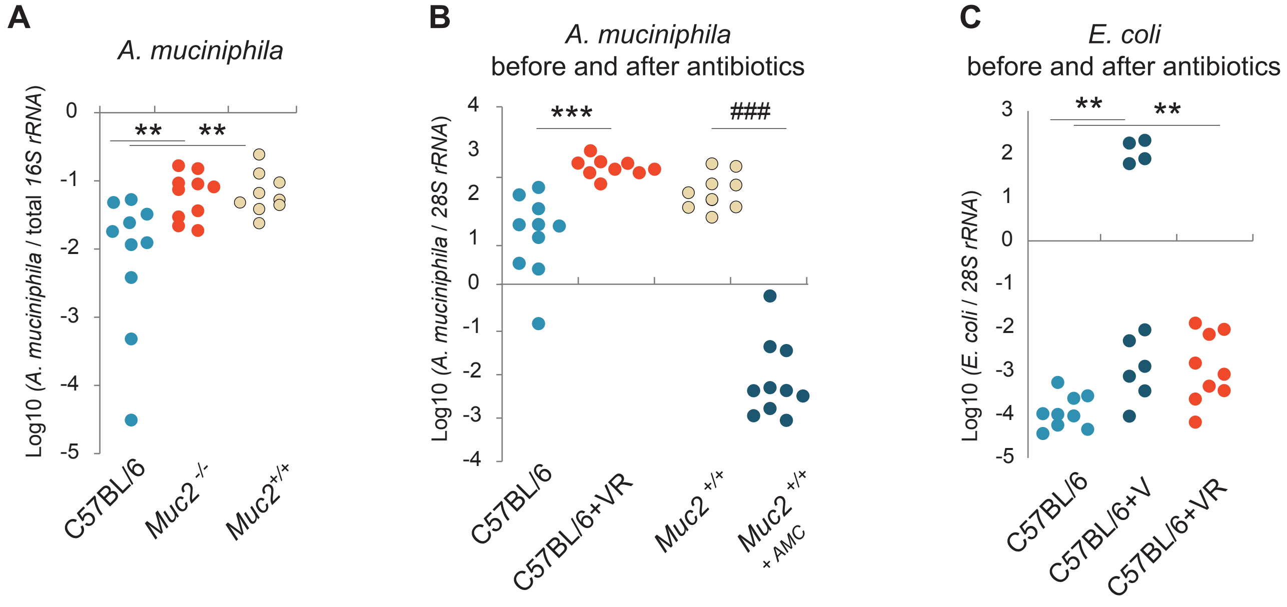
