## Supplemental table 1 for "Colitis-associated intestinal microbiota regulates brain glycine and host behavior in mice"

| Target | Primer name | Primer sequence 5' -> 3' |
| --- | --- | --- |
| *Akkermansia muciniphila* | AMUC- F | CAGCACGTGAAGGTGGGGAC |
| | AMUC-R | CCTTGCGGTTGGCTTCAGAT |
| *16S rRNA* | 16S-F | TCCTACGGGAGGCAGCAG |
| | 16S-R | ATTACCGCGGCTGCTGG |
| *28S rRNA* | 28S-F | CCTGGCGCTAAACCATTCGT |
| | 28S-R | AAAGCCCGCAGAGACAAACC |
| *E. coli* | guaB-F | TGCTTTCCGCAGCAATGGAT |
| | guaB-R | CTGCGGATCAGTCACCACAC |
| Mouse *β-tubulin (Tubb5)* | betaTub F | TGAAGCCACAGGTGGCAAGTAT |
| | betaTub R | CCAGACTGACCGAAAACGAAGT |
| *Mouse Nos2* | Nos2_F | CAGGGTCACAACTTTACAGGGA |
| | Nos2_R | CACTTCTGCTCCAAATCCAACG |
| *Mouse Il-1β* | Il1b_F | TGAAGTTGACGGACCCCAAA |
| | Il1b_R | TGATGTGCTGCTGCGAGATT |
| *Mouse Tnf-α* | Tnfa_F | CCCTCACACTCAGATCATCTTCT |
| | Tnfa_R | GGCACCACTAGTTGGTTGTCTTT |
| *Mouse Cox2* | Cox2-F | CCAGCACTTCACCCATCAGT |
| | Cox2-R | ACCCAGGTCCTCGCTTATGA |
