## Supplemental table 2 for "Colitis-associated intestinal microbiota regulates brain glycine and host behavior in mice"

### Blood metabolic profiles as revealed by nuclear magnetic resonance (NMR) spectroscopy

| Metabolite | | ***Muc2-/- vs.* C57Bl/6** | | | | ***Muc2+/+ vs*. C57Bl/6** | | | ***Muc2-/- vs. Muc2+/+*** | | |
|---|---|---|---|---|---|---|---|---|---|---|---|
| | **Test** | **p-value** | **Test** | **Z** | **p-value** | **Test** | **Z** | **p-value** | **Test** | **Z** | **p-value** |
| 2-hydroxyisovalerate | Kruskal-Wallis | 0,0372 | Mann–Whitney *u* -test | 2,418973 | 0,015565 | Mann–Whitney *u* -test | 1,436265 | 0,150928 | Mann–Whitney *u* -test | -1,36067 | 0,173618 |
| Ketoleucine | Kruskal-Wallis | 0,00001 | Mann–Whitney *u* -test | 3,74185 | 0,000183 | Mann–Whitney *u* -test | 2,45677 | 0,014020 | Mann–Whitney *u* -test | -3,43948 | 0,000583 |
| Leucine | Kruskal-Wallis | 0,0235 | Mann–Whitney *u* -test | 2,30558 | 0,021135 | Mann–Whitney *u* -test | 2,30558 | 0,021135 | Mann–Whitney *u* -test | 0,34017 | 0,733730 |
| Isobutyrate | Kruskal-Wallis | 0,0162 | Mann–Whitney *u* -test | 1,70084 | 0,088974 | Mann–Whitney *u* -test | 2,45677 | 0,014020 | Mann–Whitney *u* -test | 1,77643 | 0,075663 |
| 2-ketoisovalerate | Kruskal-Wallis | 0,0017 | Mann–Whitney *u* -test | 3,06151 | 0,002202 | Mann–Whitney *u* -test | 0,34017 | 0,733730 | Mann–Whitney *u* -test | -2,98592 | 0,002827 |
| Lactate | Kruskal-Wallis | 0,0027 | Mann–Whitney *u* -test | -0,11339 | 0,909722 | Mann–Whitney *u* -test | -2,98592 | 0,002827 | Mann–Whitney *u* -test | -2,83473 | 0,004587 |
| Acetone | Kruskal-Wallis | 0,0015 | Mann–Whitney *u* -test | 1,77643 | 0,212295 | Mann–Whitney *u* -test | 3,59066 | 0,037636 | Mann–Whitney *u* -test | 1,62525 | 0,241322 |
| Pyruvate | Kruskal-Wallis | 0,0126 | Mann–Whitney *u* -test | 2,00321 | 0,045155 | Mann–Whitney *u* -test | 2,53236 | 0,011330 | Mann–Whitney *u* -test | 1,54965 | 0,121225 |
| Acetylcarnitine | Kruskal-Wallis | 0,0131 | Mann–Whitney *u* -test | -1,02050 | 0,307490 | Mann–Whitney *u* -test | -2,75914 | 0,005796 | Mann–Whitney *u* -test | -1,92762 | 0,053903 |
| Choline | Kruskal-Wallis | 0,00001 | Mann–Whitney *u* -test | -3,36388 | 0,000769 | Mann–Whitney *u* -test | -3,74185 | 0,000183 | Mann–Whitney *u* -test | -3,06151 | 0,002202 |
| Carnitine | Kruskal-Wallis | 0,0237 | Mann–Whitney *u* -test | -2,45677 | 0,014020 | Mann–Whitney *u* -test | -2,15440 | 0,031210 | Mann–Whitney *u* -test | 0,11339 | 0,909722 |
| Betaine | Kruskal-Wallis | 0,0049 | Mann–Whitney *u* -test | -2,98592 | 0,002827 | Mann–Whitney *u* -test | -2,00321 | 0,045155 | Mann–Whitney *u* -test | 1,54965 | 0,121225 |
| Glucose | Kruskal-Wallis | 0,0471 | Mann–Whitney *u* -test | 2,07880 | 0,037636 | Mann–Whitney *u* -test | 2,07880 | 0,037636 | Mann–Whitney *u* -test | -0,26458 | 0,791337 |
| Fumarate | Kruskal-Wallis | 0,0087 | Mann–Whitney *u* -test | 1,85203 | 0,064023 | Mann–Whitney *u* -test | -1,47406 | 0,140466 | Mann–Whitney *u* -test | -2,83473 | 0,004587 |
| Histidine | Kruskal-Wallis | 0,0006 | Mann–Whitney *u* -test | 2,45677 | 0,014020 | Mann–Whitney *u* -test | 3,28829 | 0,001008 | Mann–Whitney *u* -test | 2,30558 | 0,021135 |
| Phenylalanine | Kruskal-Wallis | 0,0424 | Mann–Whitney *u* -test | 2,07880 | 0,037636 | Mann–Whitney *u* -test | 2,15440 | 0,031210 | Mann–Whitney *u* -test | 0,18898 | 0,850107 |
