## Supplemental table 3 for "Colitis-associated intestinal microbiota regulates brain glycine and host behavior in mice"

### Brain metabolic profiles as revealed by nuclear magnetic resonance (NMR) spectroscopy

| Metabolite | | ***Muc2-/- vs.*** **C57Bl/6** | | | | ***Muc2+/+ vs.*** **C57Bl/6** | | | ***Muc2-/- vs.*** **Muc2+/+** | | |
|---|---|---|---|---|---|---|---|---|---|---|---|
| | **Test** | **p-value** | **Test** | **Z** | **p-value** | **Test** | **Z** | **p-value** | **Test** | **Z** | **p-value** |
| Myo-inositol | Kruskal-Wallis | 0,028 | Mann–Whitney *u* -test | -2,08167 | 0,037374 | Mann–Whitney *u* -test | -2,19089 | 0,028460 | Mann–Whitney *u* -test | 1,27802 | 0,201244 |
| Scillo-inositol | Kruskal-Wallis | 0,0221 | Mann–Whitney *u* -test | -2,56205 | 0,010406 | Mann–Whitney *u* -test | -2,00832 | 0,044611 | Mann–Whitney *u* -test | -0,54772 | 0,583883 |
| Glycine | Kruskal-Wallis | 0,0154 | Mann–Whitney *u* -test | -2,56205 | 0,010406 | Mann–Whitney *u* -test | -2,00832 | 0,044611 | Mann–Whitney *u* -test | -1,27802 | 0,201244 |
| Inosinate | Kruskal-Wallis | 0,0122 | Mann–Whitney *u* -test | 1,44115 | 0,149542 | Mann–Whitney *u* -test | 2,55604 | 0,010588 | Mann–Whitney *u* -test | -2,19089 | 0,028460 |
